# Exploring The Behavior of Bioelectric Circuits using Evolution Heuristic Search

**DOI:** 10.1101/2022.10.23.513361

**Authors:** Hananel Hazan, Michael Levin

## Abstract

Characteristic spatial differences of cellular resting potential across tissues have been shown to act as instructive bioelectric prepatterns regulating embryonic and regenerative morphogenesis, as well as cancer suppression. Indeed, modulation of bioelectric patterns via specific ion channel-targeting drugs, channel misexpression, or optogenetics has been used to control growth and form in vitro, showing promise in regenerative medicine and synthetic bioengineering. Repair of defects, injury, and transformation requires quantitative understanding of bioelectric dynamics within tissues so that these can be modulated toward desired outcomes in organ patterning or the creation of entirely novel synthetic constructs. The major gap in the discovery of interventions for rational control of organ-level outcomes is the inability to predict large-scale bioelectric patterns - their emergence from symmetry breaking (given a set of channels expressed on the tissue) and their change as a function of time under specific bioelectrical interventions. It is thus essential to develop machine learning and other computational tools to help human scientists identify bioelectric states with desirable properties. Here, we tested the ability of a heuristic search algorithm to explore the parameter space of bio-electrical circuits by adjusting the parameters of simulated cells. We show that while bioelectrical space is not easy to search, it does contain parameter sets that encode rich and interesting patterning behaviors. We demonstrate proof of principle of using a computational search platform to identify circuits with desired properties, as a first step toward the design of machine learning tools for improved bioelectric control of growth and form.

## Introduction

Morphogenesis, or the emergence of complex anatomical shapes from groups of cells, is a critically important process for three reasons. First, it is fundamental to understanding evolution, as it implements the mapping between the genome (target of mutations) and functional body phenotypes (the subject of selection forces). Second, it is central to almost all of biomedicine: being able to control what cells build is the roadmap to definitive solutions to birth defects, traumatic injury (via regeneration), and cancer (via tissue reprogramming).^1^ Finally, it is an essential aspect of synthetic bioengineering – the efforts to build biological robots and other living constructs made to arbitrary specifications for a myriad of applications^2, 3^ Thus, it is essential to understand and quantitatively model the patterning dynamics that occur in cellular collectives.

Morphogenetic prepatterns occur via biochemical,^4^ biomechanical,^5^ and bioelectrical^6^ modalities. The latter is particularly interesting because, as in the brain,^7^ bioelectricity forms a kind of computational medium within which large-scale anatomical decisions are made by collective cell behavior.^8, 9^ Bioelectric signaling^10, 11^ includes spatial distributions of cellular transmembrane resting potentials (V_mem_) across fields of tissue (Figure 1 A), produced by the actions of ion channels and electrical synapses known as gap junctions. Such voltage gradients have now been shown to encode information about organ size, axial polarity, and various cell behaviors,^12–15^ while disorders in bioelectric signaling induced by drugs^16^ or mutation (so-called channelopathies) can trigger cancer^17–19^ and birth defects.^20, 21^ Importantly, modulation of bioelectric signaling via drugs, channel misexpression, or optogenetics has been shown to be able to induce whole organ (e.g., eye^22^) formation, regeneration of appendages under normally non-regenerative conditions,^23^ and even the formation of head structures belonging to other species.^24^ It is clear that bioelectric computations within cell groups are an important and increasingly-tractable interface through which to control large-scale growth and form.^25^

**Figure 1.**
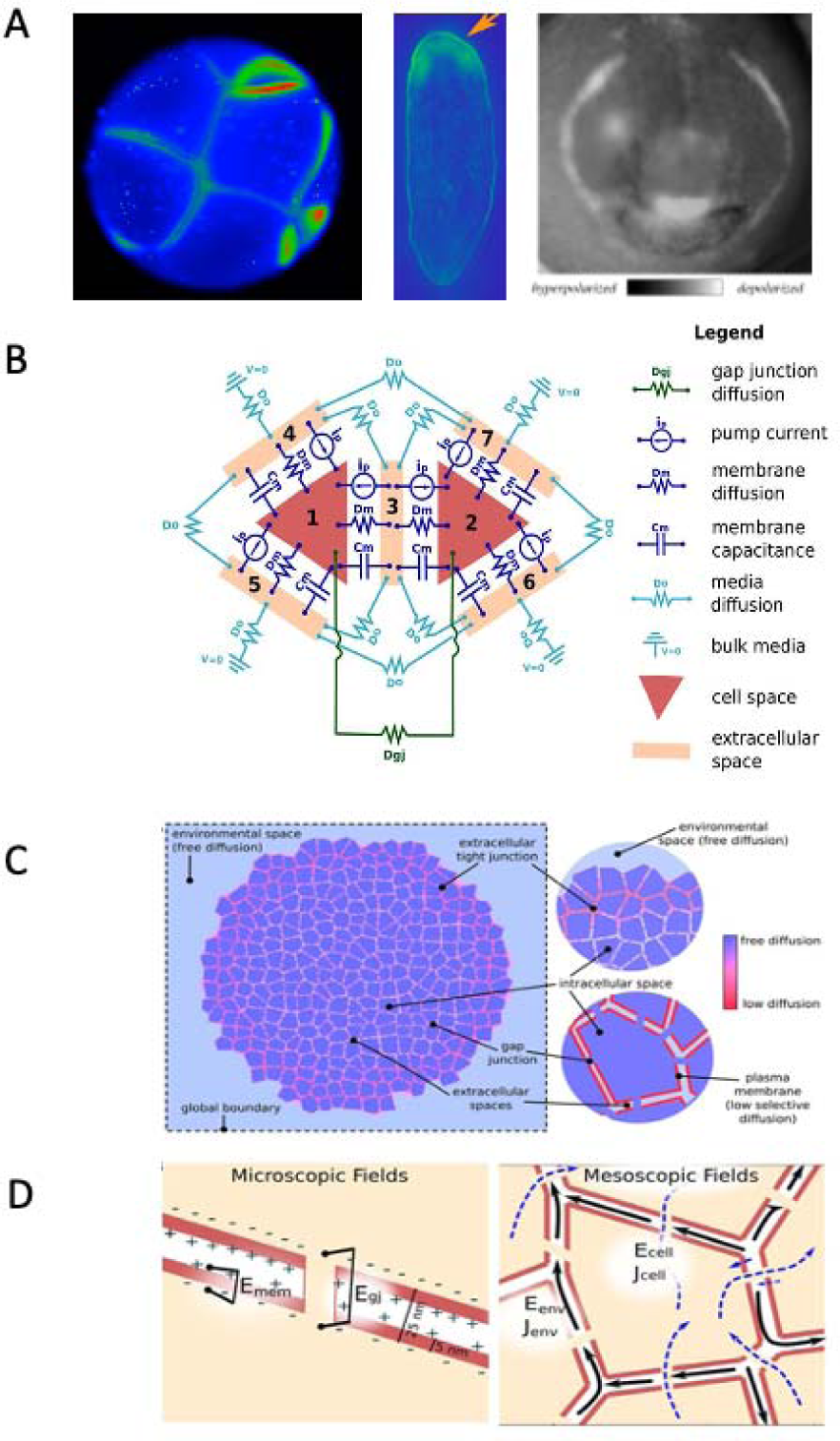
Fundamentals of developmental bioelectricity and its modeling in BETSE. (A) Sample bioelectric patterns, as revealed by voltage reporter dye technique^26^ of the cleavage-stage frog embryo, the planarian flatworm, and the developing frog face (left-to-right). Reproduced by permission from ^25^. Image of frog embryos produced by Dany S. Adams; image of planarian produced by Taisaku Nogi. (B) The fundamental “electrical circuit” implemented in BETSE, shown on a simplified geometry of two triangular cells (1 and 2) surrounded by their respective extracellular spaces (3–7). Note that in BETSE, and in contrast to the simplified image shown, cells are defined from a Voronoi diagram and are polygonal with four or more membranes, and that a larger network of 10–1000 cells is considered in simulations. Each cell–extracellular junction has a capacitive component (membrane capacitance C_m_), a “resistive” component (cell membrane diffusion coefficients, D_m_), and a variable current source (representing the action of pumps, i_p_). Transfer between two cells occurs via GJs, which are represented by a “resistive” component (D_gj_). Transfer between extracellular spaces and to the environment is handled using “resistive” components (D_o_). Boundary conditions at the global environmental boundary are represented by grounded voltage (V = 0) and fixed concentrations representing an open boundary with Dirichlet conditions. Self-capacitances for each cell and extracellular space are not shown. Image reproduced by permission from ^27^. (C) Electro Diffusive mass transport in a GJ networked cell cluster is assumed to follow three pathways. (D) Close-up view of the cell membranes. (1) transmembrane – between intra- and extracellular spaces across the plasma membrane; (2) inter-cellular – between cellular spaces via GJ; and (3) environmental – between extracellular spaces and in the global environment. Image reproduced by permission from ^27^.

One example of this kind of approach is the repair of brain defects in an amphibian model. Recent work has shown that profound defects of the brain induced by alcohol and nicotine, as well as by mutation of the critical neurogenesis gene *Notch*, could all be rescued by reinforcing the normal bioelectric prepattern that controls brain morphogenesis.^28–31^ Not only did animals regain normal brain anatomy and gene expression, but their behavioral intelligence was also returned to normal. This was accomplished by computationally modeling the ion channel circuit responsible for the bioelectric prepattern of the nascent brain, and then using that model to search for an intervention (in this case, activating the HCN2 channel) that would return the pattern back to normal. This forms a proof-of-concept for a roadmap in which bioelectric modeling is a central part of a platform that predicts electroceutical drugs for a range of biomedical indications. Importantly, the same treatment was also seen to rescue defects of the heart, face, and gut, even though bioelectric prepattern data for those organs are not yet available, reinforcing the fact that control principles may be highly conserved: cracking the bioelectric code in one context may provide actional information for interventions in others.

Thus, it becomes essential to be able to model bioelectric dynamics and to leverage those models to identify specific bioelectric perturbations (ion channel modifications using drugs or other methods). Bioelectric dynamics have been modeled using an equivalent circuit approach,^32–34^ as well as a more bio-realistic simulation using a tool known as Bioelectric Tissue Simulation Engine (BETSE).^27, 35^ BETSE is a sophisticated physiological simulator which takes as input information about tissue geometry and the presence of various ion channels and gap junctions, and reveals what the bioelectric patterns will be in that tissue as a function of time. Thus, it has the potential to reveal aspects of self-organization and symmetry-breaking,^36, 37^ which are essential to understand how complex patterns in development arise from one egg cell. It can also reveal various computations in non-neural tissue that can be exploited in the contexts of bioengineering and unconventional computing, among others.^38, 39^

However, bioelectric circuit function is complex, because voltage-gating of channels and gap junctions results in feedback loops and propagating dynamics within tissues. Even though every cell in a tissue can have the exact same channels (i.e., they all appear identical from a proteomic perspective), the channels open and close dynamically forming rich behaviors that generate order and spatial fluctuations. Thus, it is not possible to readily intuit the kinds of initial conditions or perturbations that will result in a specific outcome pattern: computational tools are needed to help scientists solve the inverse problem of predicting which kinds of ion channel properties in a field of cells will give rise to a desired V_mem_ pattern.

There are major gaps in the current ability to predict bioelectric patterns and their temporal evolution in tissue. Moreover, it is not known what kinds of patterns typical bioelectric circuits can form – what are the possible behaviors to be found in the space of all possible bioelectric circuits? While chemical reaction-diffusion systems have been studied extensively, it is not clear what the computational and functional capabilities of bioelectric circuits are. To address these questions, here we undertake two main aims. First, we begin the development of tools for the bioinformatics of form,^40, 41^ using techniques from machine learning to help discover conditions for bioengineering/biomedical settings. Second, we start mapping out the phenotypic bioelectrical space to construct an ontology of the kinds of patterns that bioelectric circuits can establish, facilitating the study of the emergence of complex bioelectric behaviors from a homogenous initial state.

### Approach

Specifically, we seek to solve the inverse problem: given a (disease, or initial) state with bioelectric pattern P’, and a known correct bioelectric pattern P, which native ion channel proteins should be opened and closed in order to convert P’ into P? Once this is known, existing ion channel modulation techniques can be selected that would perform the needed manipulation of the ion channels. Using BETSE, any bioelectric circuit can be simulated through time.^27, 35, 42^ However, it is very computationally intensive, and thus is not suited for exhaustive search. We produced a heuristic search algorithm tool to test one specific hypothesis - that distinct BETSE circuits exist, and can be found via the proposed ML approach, and that those circuits have several desirable properties: (1) self-organization, (2) robustness against transient (external) induced bioelectric perturbations, and (3) memory of this pattern that can be reset (permanently altered) to a different robust pattern by a specific bioelectric input stimulus. Such circuits would be important as testable models for building novel synthetic biological constructs such as biobots,^43^ and as targets for design of novel synthetic bioelectrically-controlled tissues.^44–46^

Within BETSE, cells are represented as shown in Figure 1 B, and simulations can be done of multicellular tissues that express specific complements of ion channels and pumps, with specific properties. Moreover, BETSE allow us to use external or internal interventions (mimicking drug or optogenetic stimuli) that interact with the ion channels, extra cellular substances, voltage interventions or gap junction of the cells in the simulated tissue (see Figure 1 C).

## Methods

The work done in this paper uses two different programs: the tissue simulation program and the heuristic search program. BETSE (BioElectric Tissue Simulation Engine, https://github.com/betsee/betse) is an open-source cross-platform discrete exterior calculus simulator for 2D computational Multiphysics problems in life sciences (developed by Alexis Pietak at the Allen Discovery Center at Tufts). BETSE simulates the interaction of cells in a tissue and takes into account the electro-diffusion, electro-osmosis, galvanotaxis, voltage-gated ion channels, gene regulatory networks, and biochemical reaction networks (e.g., metabolism). Our heuristic search uses our own flavor of genetic algorithms that are tuned specifically to run efficiently on High-Performance-Computer (HPC) clusters (see Figure 2 A and B) to find the relevant parameter for our tasks.

**Figure 2:**
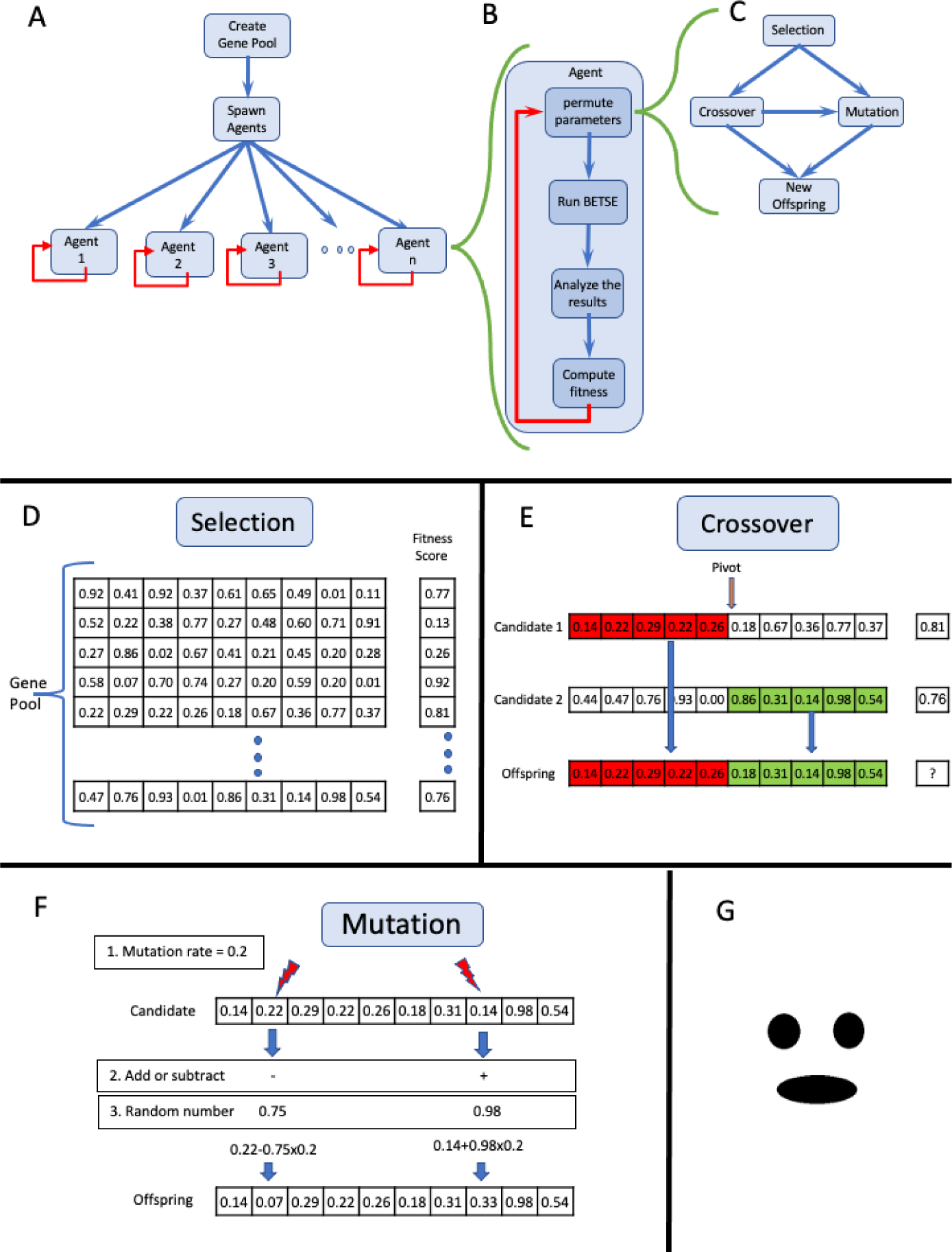
Schematic of the heuristic search algorithm. (A) On initiation, the main algorithm creates the gene pool by choosing a random set of parameters and then spawns the agents. (B) Each agent is an independent process that performs an endless loop of selecting a new gene, running BETSE, and evaluating the fitness score of the result and starting over. (C) Each agent generates a new offspring by following this diagram. The agent selects the candidates from the gene pool, performing either Mutation or Crossover and then Mutation on the new offspring. (D) The Selection operation, *select the candidates from the gene pool using a weighted randomness base on the normalized population fitness score*. (E) The Crossover operation uses parts from the candidates’ chromosomes from the last step to create a new chromosome. In this example, we randomly choose a pivot location and combine the first candidate chromosome from the left side of the pivot and the chromosome from the second candidate right side of the pivot to create the new offspring. (F) The Mutation operation gets from the last step (Selection or Mutation) one chromosome and determines how many mutations to execute on the given chromosome, along with the strength of the mutation (mutation rate). In this example, the Mutation operation executes two mutations that change two genes from the offspring chromosome. (G) The fitness function can use a mask file for evaluating the tissue pattern. This mask file is an image file that contains black and white pixels. The fitness function uses this mask as a set of coordinates of the black/white pixels to pinpoint the exact cells in the tissue to evaluate. For example, the fitness function averages the cells V_mem_ located under the black pixels and compares it to the average V_mem_ of the cells in the white areas. The bigger the difference the higher the fitness score.

A genetic algorithm is a heuristic stochastic exploration method that uses partial information to optimize a desired fitness function. The algorithm takes inspiration from the Darwinian natural selection process and uses similar operators – such as Selection, Mutation, and Crossover - to explore a large space of possibilities. Here, the genes in the genetic algorithm serve as the parameters for the experimental configuration test of the BETSE simulator. The genetic algorithm performs parameter tuning of the gene based on our fitness function, which we set for each task below. The genetic algorithm is composed of multiple agents that share a gene pool, run simultaneously, and tune asynchronously.

The genetic algorithm works as follows: its main program starts by creating the gene pool by choosing a random set of parameters, and then spawns the agents. Each agent is an independent process that performs an endless loop: selecting a new gene, running BETSE, and evaluating the result. Each looping agent process starts by selecting sets of genes from the gene pool, performing Crossover and/or Mutation, and thereby creating a new set of genes (offspring). Then the agent evaluates the offspring by running BETSE, and then runs the fitness function against the BETSE results. Each agent evaluates whether its resulting gene fitness score is better than any other gene in the population. Depending on the fitness score, if any one of the genes in the population ranks lower than the agent’s fitness score, the agent replaces the lowest fitness score with its own, thus improving the fitness score of the general population.

### Genes and Population

The population (gene pool, see Figure 2 D) contains all the genes in the population and their fitness score. The gene pool is shared between all agents and is used by the selection process in each running agent. Each genome in the population contains a set of 33 genes corresponding to the parameters of our task. Each one of the 33 genes specifies a number between 0 to 1, and the metadata on each gene is stored separately. The metadata of the genes is the effective range of each parameter, as well as some other properties relevant to the parameter (e.g., “round 19” means that genes that are represented by a 64-bit floating point number will be rounded to 19 digits). The genetic search algorithm tries to explore this 33-dimensional space to maximize the fitness score of the given task.

### Genetic Algorithm Steps

The genetic algorithm uses genes to represent the parameters in the system. To operate on those genes the algorithm employs operators similar to those found in Darwinian natural selection. In natural selection, the population represents the most successful candidates of the previous generation, and the next generation is created from the candidates with the preferred traits. The genetic algorithm (see Figure 2 C) operates as follows: To create the new offspring, it first needs to select a couple of gene strains from the gene pool. Depending on the diversity of the gene pool, the algorithm selects either the Crossover operation or the Mutation operation.If the diversity of the gene pool is low, the chances of choosing Crossover are reduced, since crossover between similar genes does not contribute much to the guided search. In any case, if Crossover has been selected, there is still a 50% chance that mutation can occur on the gene that results from the Crossover operation.

### Selection

The first step of the agent is to select parents to create offspring. The agent can choose one, two, three, or four candidates to serve as a baseline from which to create the offspring. The Selection process uses weighted random selection to choose suitable candidates from the population (see Figure 2 D) based on their fitness scores. This method tends to increase the percentage of “better-fit” genes in the gene pool. The weighted random formula uses a normalized population fitness score according to the following pseudo-code:

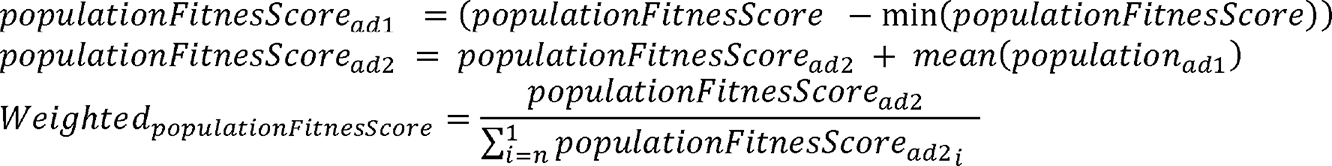

The weighted score of the population is used as a discrete probability distribution for the random function that chooses the candidates for the next operation (Crossover or Mutation).

### Crossover

Given the parents’ gene vectors from the Selection process, the Crossover process merges random parts of each parent to create the offspring vector. Crossover starts by randomly selecting (via uniform distribution) a set of pivot points where the vector merge starts (see Figure 2 E). The Crossover operation can use two or more candidates in order to create offspring. The number of pivot points that can be chosen by the Crossover operation can vary from *1* to *gene length – 1*, and will depend on the mutation rate, a variable set by the user. The result of this operation is a strain of gene that shares many parts with its parents, each parent having contributed different gene strains with their own unique features.

### Mutation

The input for the Mutation operation can come from the Selection operation or the Crossover operation. In either case, depending on the mutation rate, the Mutation algorithm chooses one or more genes to be mutated. Each chosen location is added or subtracted from a random number between 0 to 1 and multiplied by the mutation rate variable (Figure 2F).

### BETSE parameters and initialization

The total parameter space of our heuristic search was 33 parameters; 18 parameters relate to the cell’s properties and 15 parameters relate to the initiation of the environment of the tissue (for the list of the parameters see the Supplementary Materials at https://dataverse.harvard.edu/dataset.xhtml?persistentId=doi:10.7910/DVN/U1QRS8). Of the 18 parameters for cell properties, 11 are common to all permutations, and 7 are related only to voltage dependent ion-channels. In the first iteration of the BETSE simulation, all the cells have the same starting point, meaning they all have the same V_mem_; therefore, no diffusion can occur since there is no voltage gradient present in the tissue. To address this issue, we introduce only in the first step of the simulation a symmetry break that introduces gradients in the tissue, and that then disappears right after the first step. This allows the diffusion reaction between the cells to begin. In our results presented here, 15 of the 33 parameters relate to the BETSE initializing the environment, and those are used for symmetry breaking at the beginning of the BETSE run. Each symmetry breaking point is a circular intervention that uses three parameters: one for the coordinate, one related to the radius, and one related to the strength of the intervention. In total, we use five symmetry breaking points that correspond to those 15 genes.

### Fitness function

As part of the heuristic search algorithm, the result of each BETSE simulation needs to be evaluated to see how well it fits a desired profile. The evaluation process returns a score that represents whether our current result is closer to or farther from our desired pattern. For example, searching for a cell tissue that shows a specific V_mem_ pattern resembling a “smile” image, we first provided an image with the desired pattern (Figure 2 G). The fitness function compares the V_mem_ of the cells that are located in the marked (black) pixels, and the V_mem_ of the cells that are located in the unmarked (white) pixels of the mask. Thus, the fitness algorithm counts how many cells located in the desired smiley image (marked black in the fitness mask) have a higher value compared to cells that are outside of the mark. The fitness score counts the number of cells that are located in the white pixels and divides them by the total of cells in the unmasked image. The total fitness score that is returned to the genetic algorithm always ranges from 0 to 1.

## Results

To use BETSE, one needs to set many parameters; these are related to cell morphology (cell properties such as cell membranes and ion-channels), and to environmental properties (such as chemicals that exist in the bath, temperature of the environment, etc.).Some of the parameters are easy to manipulate and control in the real world, and some are not. We fixed the non-negotiable parameters (e.g., materials properties of the ions that affect diffusion etc.), and let the heuristic search algorithm search the space of those parameters that experimentalists can readily control.

All BETSE simulations start in a homogenous state where all cells in the tissue have the same values. In this situation, diffusion cannot drive the system from this hard stationary point; therefore, we included physiological noise (stochastic perturbations during the initialization phase of the model) to enable spontaneous self-organization. We chose nine specific tasks of biological relevance, in which to examine the capabilities of the heuristic search algorithm to find the right parameters in the multidimensional bioelectrical space. The goal of the heuristic algorithm was to propose a bioelectric circuit that fits our desired task. The tasks given to the heuristic search ranged in complexity from finding a simple bio-electrical circuit to finding complex bio-electrical circuitry. The results presented here use different shapes of tissues: tasks 1-4 were made on a square tissue, while task 5 used a circular tissue, task 6 used many different shapes of tissue, and tasks 7-10 used an oval tissue resembling a planarian flatworm. Moreover, to better understand the interplay between transcriptional and bioelectric mechanisms, we also used two different gene-regulatory network (GRN) mechanisms for our configurations for tasks 1 and 5 (as described in the Supplemental BETSE config files). The different GRN mechanisms added some complexity to the V_mem_ patterns, but the search for the other parameters was the same as in the prior simulations. All the configuration details used in this paper can be found in the Supplementary Materials. The results below are representative examples from the heuristic search; the results for tasks 1-9 are the best fit for our fitness function. Examples in Supplemental Figures 1-23 show a sampling of interesting patterns employed in the process of finding the best fit for task 5.

### Task 1: Tissue with as little V_mem_ changes as possible over time, and cells maintain homogenous V_mem_

We first sought to find a set of initial conditions that create a stable homogenous field – corresponding to an ion channel configuration that can support a stable, adult tissue region which is meant to be isopotential and remain constant for a long period of time. Thus, we searched parameter space for V_mem_ outcomes with a very slow, almost zero activity. Moreover, the cells’ activity is characterized by slow change throughout all cells, meaning the variance between cells will be close to zero. The heuristic search explored the parameter space for a set of parameters that return the highest score. The fitness function was set to reward a tissue that has the lower V_mem_ variance between the cells. Each run of the heuristic searches for the genetic algorithm homing in on one solution that maximizes the fitness score. We found many such solutions, with the same voltage pattern and temporal characteristics but with different parameters. The example presented in Figure 3 is one of the examples we found, using the parameters from the configuration files in Supplement 1 (file directory Task 1). We conclude that bioelectric parameter space readily produces circuits with stable, homogenous bioelectric prepatterns.

**Figure 3:**
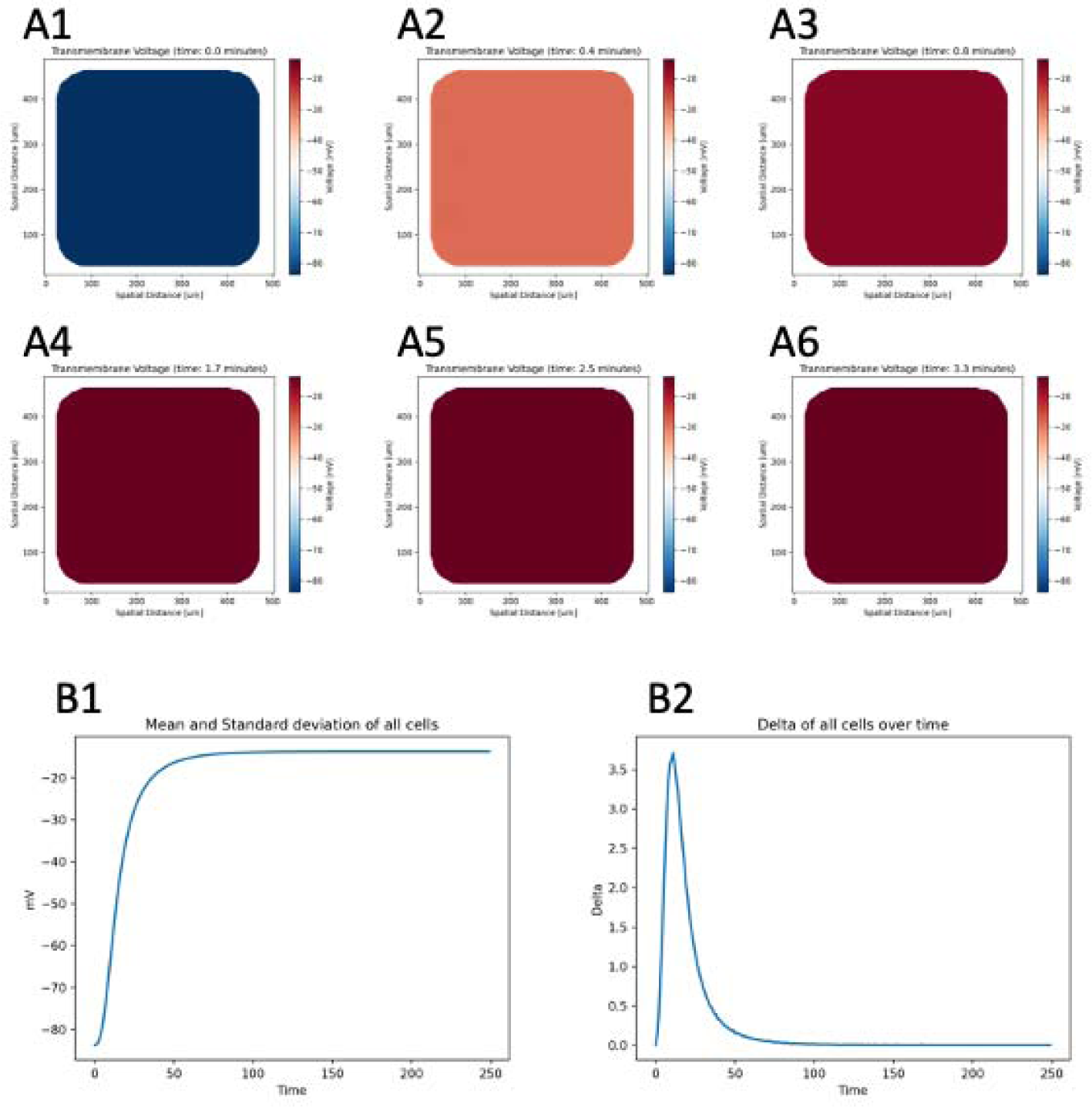
Results of Task 1, A tissue that has as little V_mem_ change as possible over time, and the cells maintain homogeneous V_mem_. Here is a temporal snapshot of a configuration that creates a cells’ tissue from initiation (A1) to the stable point (A3) 48 seconds from initiation to the (A6) end of our simulation at 3.3 minutes. (B1) is a timeline of average V_mem_ of cells throughout running of the simulation. The tissue starts with an average of -80mV and stabilizes at about 50 seconds to a V_mem_ that stays constant throughout the experiment. Throughout the experiment the variance between cells was negligible. (B2) shows the delta between cells over time.

### Task 2: Tissues that have as little change as possible through time, but with high variance between cells

We next searched for a circuit that would set up some rich spatial prepattern (regardless of what it might be) and maintained it stably over time. This case corresponds to a developmental scenario in which a patterned regionalization needs to be established (e.g., organ compartments such as in the bioelectric face prepattern) and then maintained for significant time. The heuristic search found many configurations that could fit this description. One of these is shown in Figure 4, where the tissue immediately stabilizes on a certain pattern and does not change that pattern throughout the experiment running time. This example uses the parameters from the configuration files in Supplement 1 (directory Task 2).

**Figure 4.**
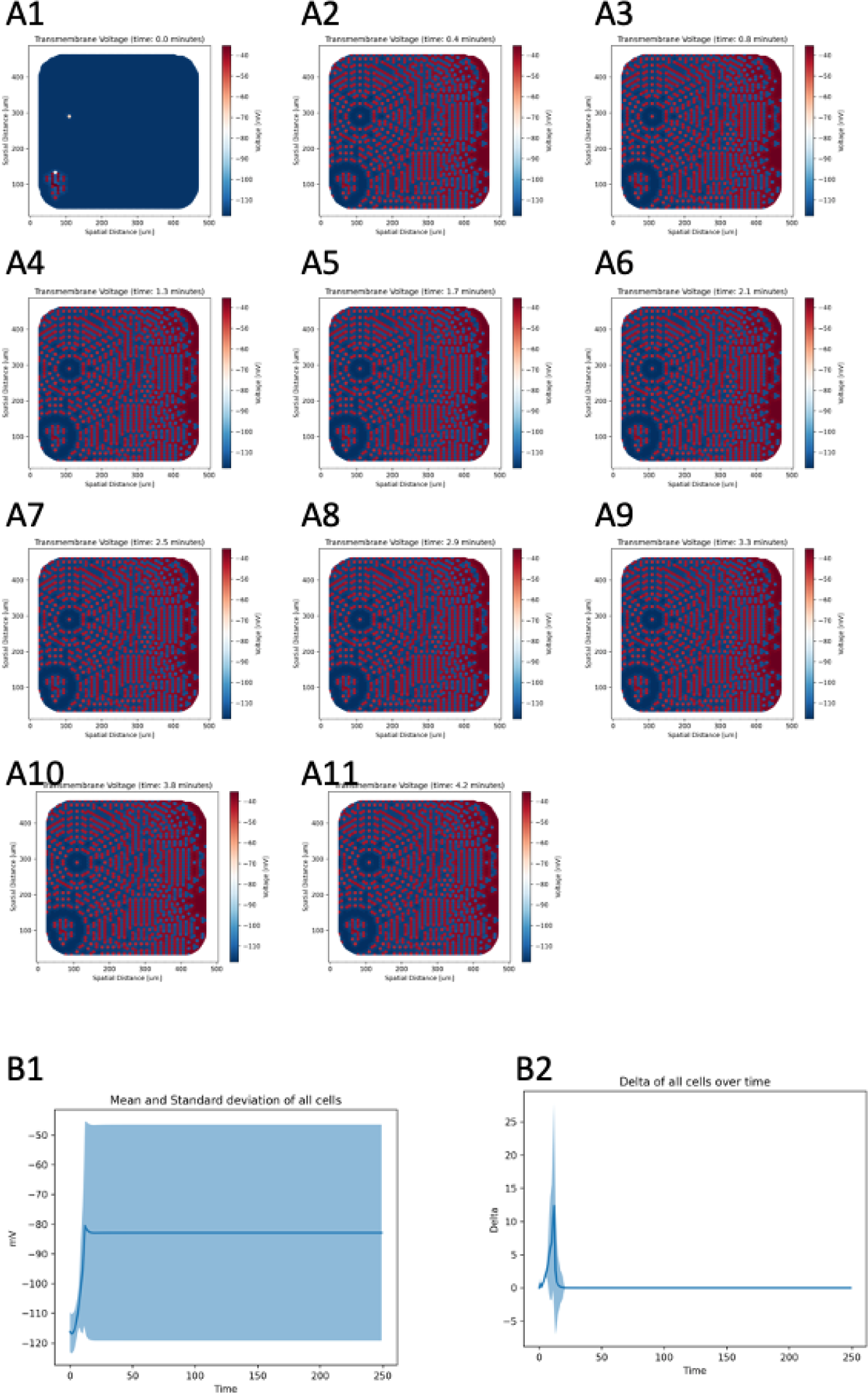
Task 2: Tissues that have as little change as possible through time, but with high variance between cells. Temporal snapshots of configuration of cells tissue from (A1) initiation depicting a tissue with a pattern that shows high variance at the immediate (A2) stabilization point and continuing to the end (A11) of the experiment. (B1) is a timeline of the average of cells Vmem throughout running of the simulation. The tissue starts with an average of about -117mV and stabilizes at about 15 second to a Vmem that then stays constant throughout the experiment. Throughout the experiment the variance between cells was ignored. (B2) shows the change in Vmem level between consecutive frames over time. It is seen that there was high change in the beginning of the run, but zero change for the remainder of the run.

### Task 3: Fits a specific V_mem_

We next simulated a scenario where, in the context of regenerative medicine or bioengineering, one wants a circuit that stabilizes at a particular V_mem_ value, in order to (for example) normalize a depolarized tumor or kickstart regenerative response by depolarizing mature tissue. In this example, we set the fitness function to give the maximum score to a tissue where the V_mem_ of all the cells in the tissue was stable at -35mV, with no variance among the cells. The heuristic algorithm found many configurations that fit our description, one of which is shown in Figure 5 A.

**Figure 5.**
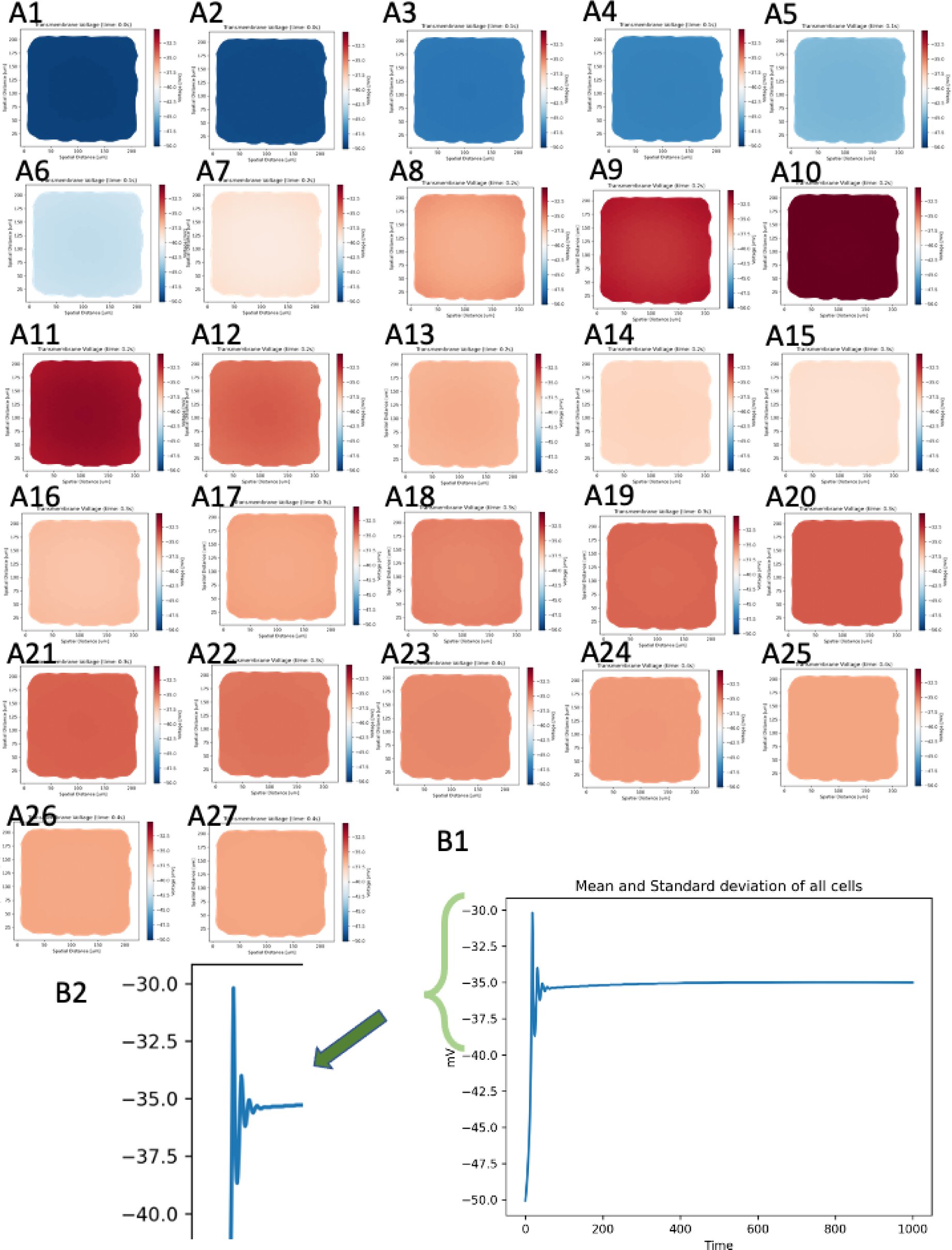
Task 3, Tissue that fits a specific V_mem_. Temporal snapshots of the cells tissue from initiation (A1) to the stable point (A24) at 0.5 seconds. (B1) The tissue starts with an average of -50 mV and overshoots and undershoots the desired V_mem_ until stabilization at about 0.5 seconds. (B2) For easier viewing, we enlarged the stabilization process that occurred in the start of the run.

We then investigated how the tissue we found would behave under perturbation, and whether it would then be able to recover its desired V_mem_; we need to understand the robustness of such circuits, because in application it is important to know if target cells will bounce back from desired interventions or environmental noise. Thus, we implemented an external stimulation that was intended to shift the V_mem_ away from our desired stable V_mem_. The external stimulation temporarily changed membrane permeability to Na^+^ in all the cells’ tissue, corresponding to a transient opening of sodium channels. We found a range of responses to the stimulation. 1. In some of the tissues found by the heuristic search, stimulation caused the tissue to destabilize away from the desired V_mem_. 2. In some tests, the tissues just ignored the stimulation and maintained the desired V_mem_. 3. In some tests the tissues almost immediately stabilized on the desired V_mem_. 4. In some tests (see Figure 5B) the results were similar to the behavior of a PID controller,^47^ where the tissue actively tried to stabilize on our desired V_mem_ despite the stimulation. The tissue that behaved similarly to PID controllers showed a strong stable point in stabilization on the desired target, and the simulated cells quickly converged to the desired V_mem_ despite the stimulation. From the many examples we got from the heuristic search, we chose to show here one interesting example of the tissue that had homeostatic behavior at the cellular level, using parameters from the configuration files in Supplement 1 (directory Task 3). Such behavior is not only useful for understanding resistance of cells to bioelectrical modulation (e.g., cancer cells during normalization attempts), but it is also a desirable property for synthetic biology constructs exploiting bioelectricity.

Note that we did not need to have the fitness function specifically reward for stability: the homeostatic property was an emergent feature akin to a “free lunch” (in the physics sense) in some of these circuits. This could have interesting implications for the evolution of patterning mechanisms that involve bioelectric dynamics.^48^

### Task 4. Tissues with rich spatial structure that also change through time

We next pursued a variant of task 2, using parameters that produced a tissue that was not only spatially regionalized, but was also changing actively as a function of time. This enabled us to ask whether it was possible to establish very small spatial domains (i.e., cells close to each other that had very different voltage levels), and also to create highly active tissues where the bioelectric pattern did not have the stable character we observed in prior tasks: could the temporal and spatial domains of the bioelectric prepattern be similarly active? We set the fitness function to return a maximum score for a tissue that showed high variance between V_mem_ of the neighboring cells, and to return a high score for high variance in its own V_mem_ value throughout time. With this setup, the genetic algorithm found a set of cell parameters that produced much variance between cells. Moreover, the variance between cells was high not only spatially, but also temporally. In Figure 6 it is seen that the variance of the cell throughout time is high and fluctuates; the delta between frames is high; and the variance of frames is also high (see parameters from the configuration files in Supplement 1 (directory Task 4)). We concluded that the space of possible circuits contains those with dynamic spatial evolution, as well as the stable ones shown in prior tasks.

**Figure 6.**
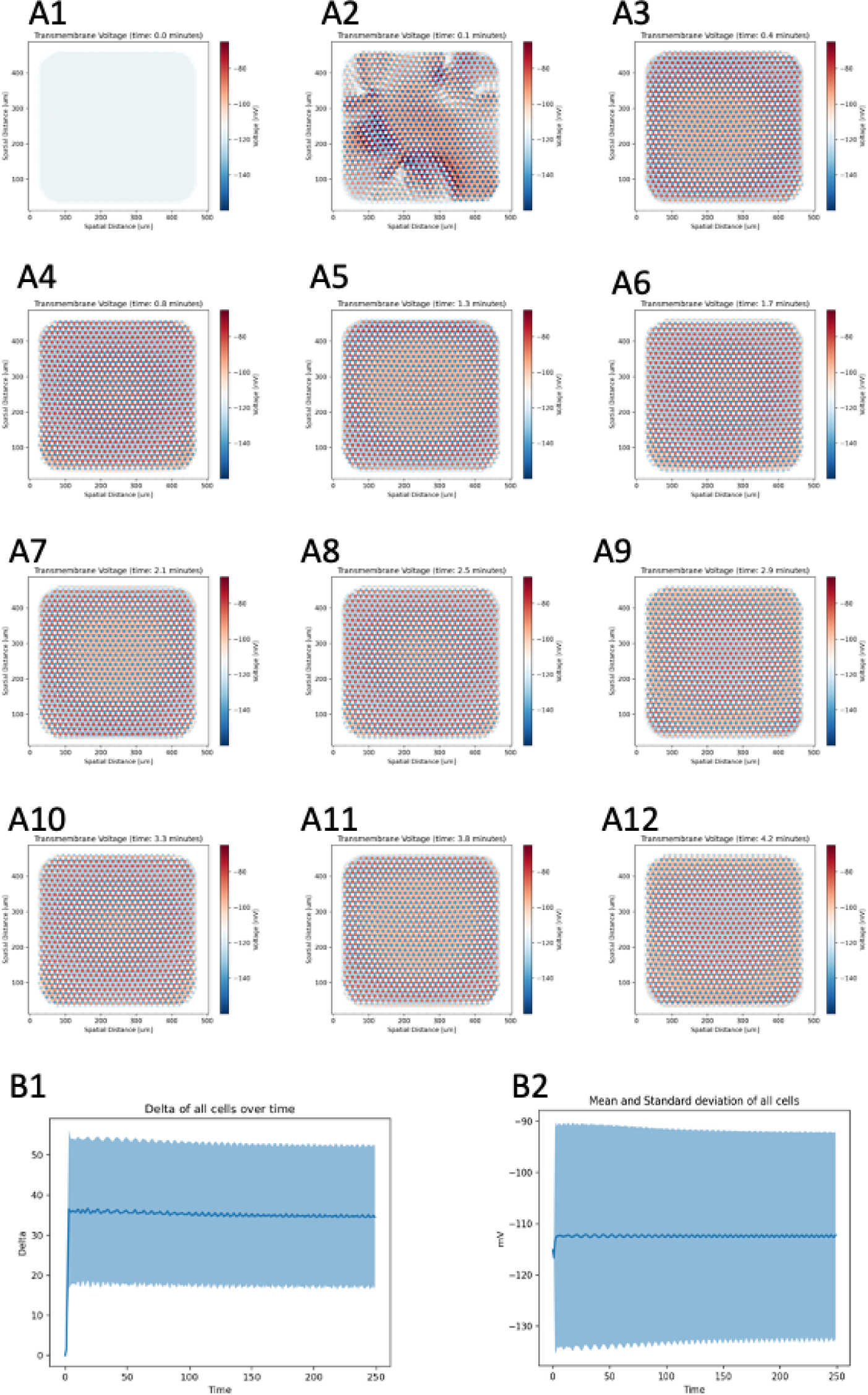
Task 4: A tissue that has high variance of V_mem_ between cells, in both spatial and temporal domain. Temporal snapshots of tissue activity show high variance between neighboring cells and across time. The timeline starts with subfigure (A1) and ends with (A12). (B1) shows the delta between consecutive frames over time. (B2) shows the average of all the cells V_mem_ throughout time, and standard deviation of the cells V_mem_ throughout time. Marked change occured between cells (B1) and between cells in time (B2) throughout the experiment.

### Task 5: Tissue that is stable on a specific pattern

One of the key features of embryonic development and regenerative repair is its robust ability to reach a species-specific pattern. Likewise, bioengineers will want to be able to express channel complements in target cells that will enable them to autonomously establish an arbitrary, pre-defined pattern. Thus, we next explored the parameter space for a pattern that features concentric bands of V_mem_ – a “bullseye”. The fitness function gave a high score to tissue that exhibited features similar to those in the desired pattern shown in Figure 7 A1. The fitness function scores high when the V_mem_ between different areas in the tissue behave similarly to the Figure 7 (A1), but not focusing on the exact V_mem_ values rather on the delta between different areas (in line with the observation that bioelectric patterns are interpreted by tissue as differences, not absolute values ^29^). Thus, the desired values of the cell V_mem_ should be highest in the yellow circle, and gradually lower in the outer circle. The heuristic search found a proximity to the desired pattern, although not exactly a perfect match (Figure 7 B1 to B6). This example uses the parameters from the configuration files in supplement 1 (directory Task 5 - bullseye).

**Figure 7.**
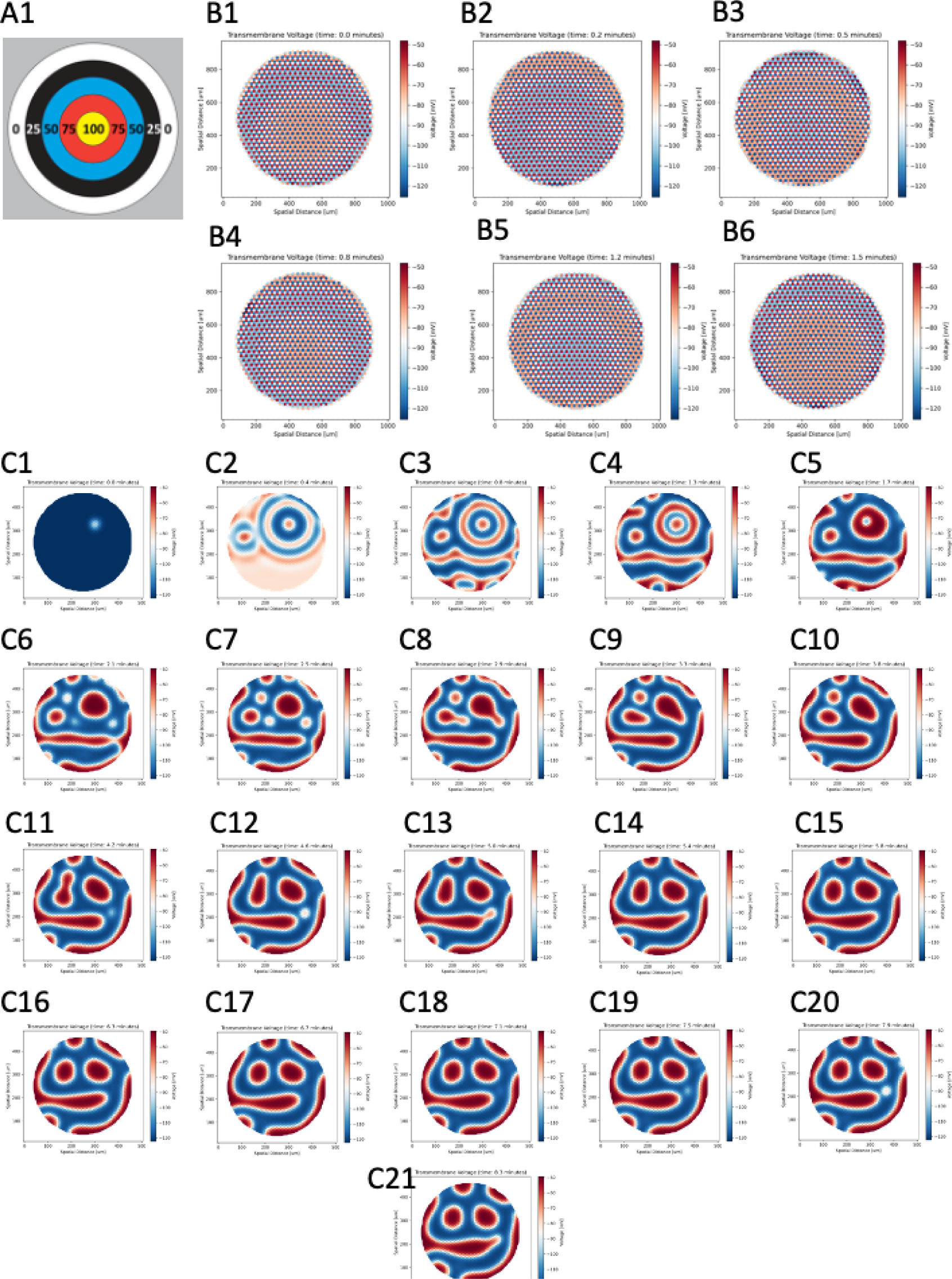
Task 5: Tissue that is stable on A specific pattern. A1) An example for a specific pattern – the bullseye. The fitness function gives a high score for tissue that shows a similar V_mem_ pattern to the bullseye image. The number in the rings of the bullseye represents the percentages of the V_mem_ value. The highest V_mem_ value of the tissue is in the center (yellow) and the lowest is in the outer ring (white). (B1-B6) A snapshot of the cell tissue activity shows the similarity to the bullseye throughout time (C1-C21) Temporal snapshots of the tissue V_mem_ activity that progresses in time toward the Smiley target. The tissue started with the symmetry breaking (C1) and progressed toward the desired pattern that appears in (C14), the smiley pattern maintains until the end of the simulation (C21).

We then searched parameter space for a circuit that would establish a V_mem_ pattern matching the “Smiley face” pattern shown in Figure 2 G, because this was structurally similar to the bioelectric face prepattern shown in Figure 1 A. The search found a configuration of tissue that started with V_mem_ patterns as result of our symmetry breaking (Figure 7 C1) and slowly morphed toward the desired Smiley face (Figure 7 C14) near the end of our simulation (for the full timeline see Figure 7 (C1-21)). This example uses the parameters from the configuration files in Supplement 1 (directory Task 5 Smiley). We conclude that it is possible to search for specific patterns; but using computational constraints, it is not easy to find a channel configuration for precisely the desired pattern.

### Task 6: Patterns with insensitivity to shape or size of tissue

An important aspect of some applications, especially in vivo, is the case where the cell field does not have a simple desired shape – for example, because morphogenesis alters the tissue geometry, or because a wound bed may have complex and variable shape. Thus, we asked how much the patterns we see depend on the shape of the tissue in which they were originally discovered. We took two of the interesting configurations we got from the heuristic search in prior experiments and checked whether the pattern found can be maintained with different tissue shapes. We tested two different ion channel complements; one configuration features continuous (spontaneous) waves of V_mem_ excitation, and the other configuration features stable high and low V_mem_ patterns. We tested whether the patterns observed in the original shape would also arise if we changed the shapes of the tissue in way that was not present during the search process. We also examined stability with respect to cell number, as one of the key remarkable features of developmental biology is the ability to produce the same morphological pattern using radically different numbers of cells.^49, 50^

In the next set of experiments, we simulated tissues in the shapes of a circle, snake, star, heart, flatworm, and ellipse, with 0.5X, 2X, and 3X the original number of cells. The results shown in Figure 8 reveal that the main features of the tissue activity are maintained regardless of the tissue shape and cell count. Figure 8 (C1-C6) shows our finding that the tissue with the slowly-changing pattern maintained its ability to generate a similar pattern with the same characteristic of low and high V_mem_. The exact voltage image was not maintained, but the overall pattern was very similar across shapes. Similarly, in Figure 8 (A1-A4, B1, B4) the tissue with high activity of excitation maintained the excitation feature and had similar refractory periods; however, different shapes and cell counts altered the frequency of the excitation waves by a small amount as a result of feedback from the cells in the, the time it took for one wave to reach the end of the tissue, and the interference between waves as a result of tissue size. Our conclusion is that some characteristics – such as frequency of activity, high and low distributions and duty cycles of the V_mem_ – are sensitive to the size and shape of the tissue. However, the main characteristic of the tissue, such as stripes, variance between cells, and ability to have action potential, all are maintained regardless of tissue size and shape.

**Figure 8.**
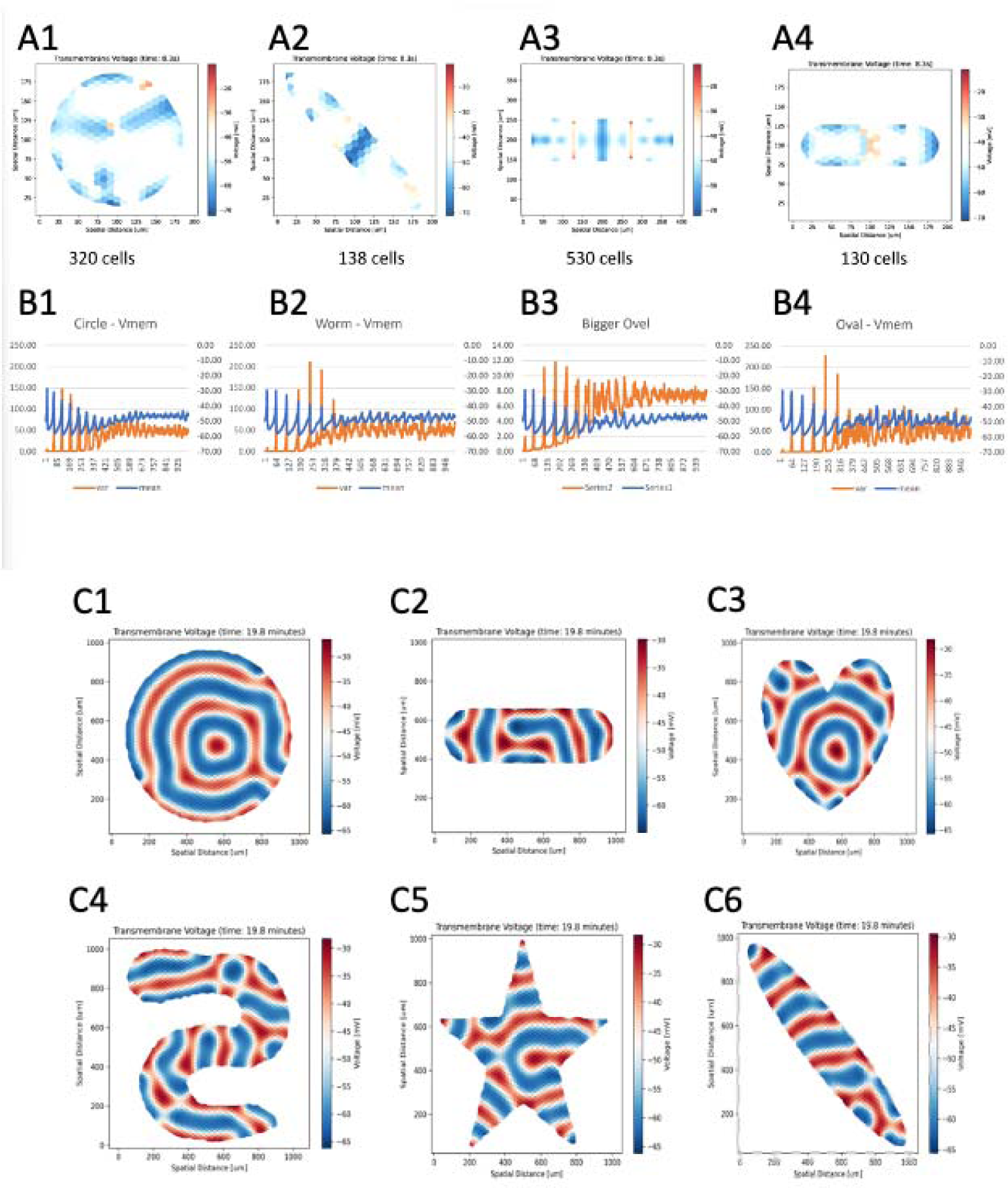
Task 6: Patterns with insensitivity to shape or size of tissue. Similar excitation patterns appear in different sizes or shapes of a tissue. Here we show the mean and variance of the cells for Circular tissue (A1, B1), Worm-shaped tissue (A2, B2), Elliptical tissue with 530 cells (A3, B3), and Elliptical tissue with 130 cells (A4, B4). The bio-electrical configuration file shows the same V_mem_ features in different tissue shapes and cell count: circle (C1), ellipse (C2), heart (C3), snake (C4), star (C5), and worm (C6). The tissue maintains the same characteristic of activity regardless of shape or number of cells.

### Task 7: Tissue that self-heals after receiving a stimulus

Regeneration and regulative development are common in biological systems;^51^ likewise, it is a goal of biorobotics research to make living machines that self-repair. To help understand and re-create bioelectric patterns that can heal after injury, we searched for a circuit that could heal its stable pattern after a perturbation.^52, 53^ In this experiment, the fitness function checked whether the V_mem_ of the tissue returned to its resting state after intervention. The virtual flatworm tissue in Figure 9 A was divided into several regions, plotting the average V_mem_ of each region. We found a tissue that started by stabilizing on its resting voltage, but then when the external stimulation started, it temporarily changed the membrane permeability of all its cells. The fitness function scored a tissue that shows any reaction to the stimulation; the highest scores was given to tissue that returned to the resting state when the stimulation was over. The genetic algorithm found a configuration that created a cell tissue that self-healed its bioelectric pattern after intervention (see Figure 9 B). This example used the parameters from the configuration files in Supplement 1 (directory Task 7), and showed that it is possible to identify homeostatic multicellular bioelectric patterns.

**Figure 9.**
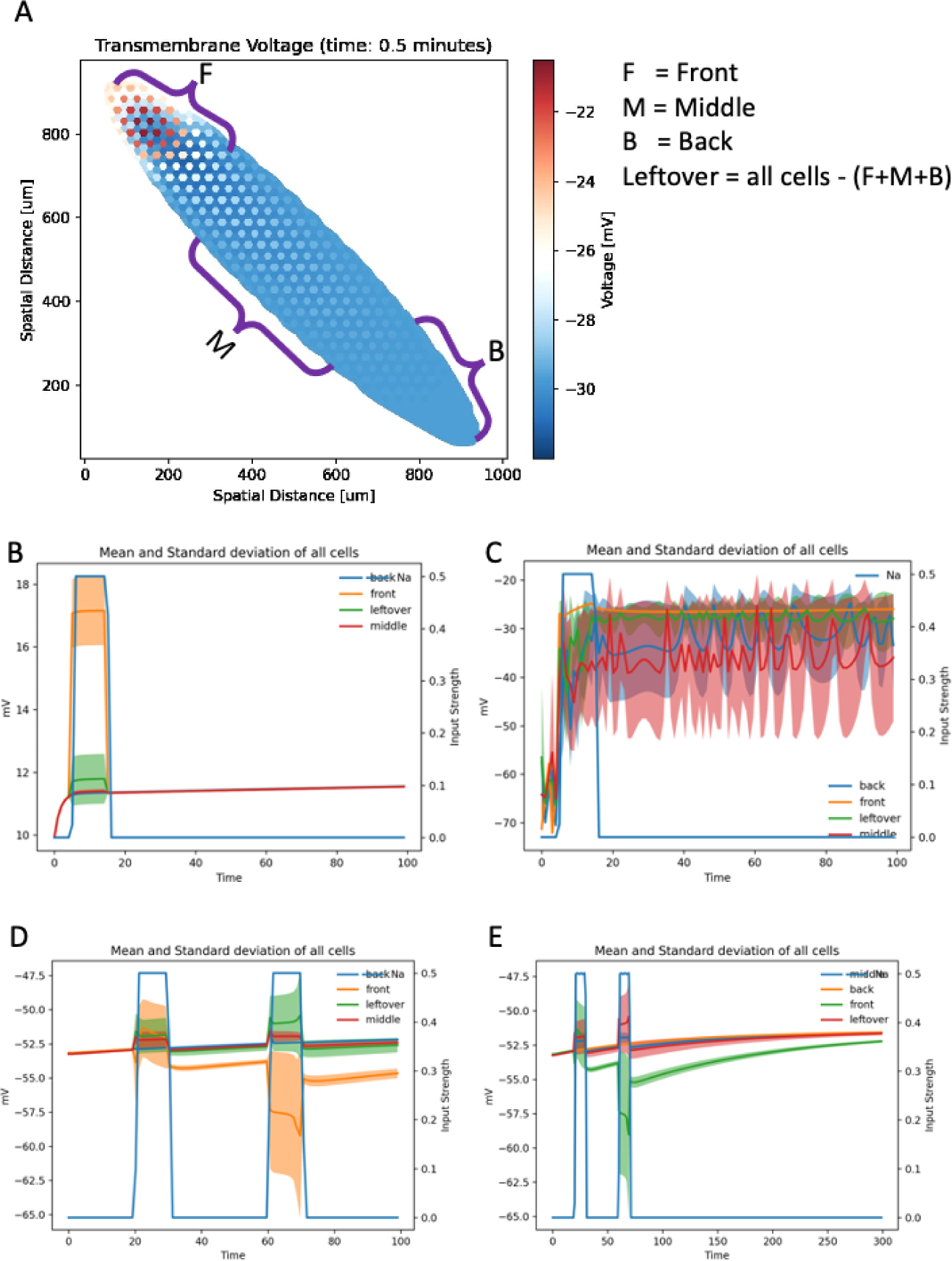
Tissue stimulations in tasks 7,8 and 9, A) Worm-shaped tissue was divided into Front, Middle and Back regions. Because those three parts did not include all the cells in the worm, we designated a fourth section, called Leftover, that contained all cells not included in the first three. We used those regions in the following tasks. The stimulus in Tasks 7, 8, and 9 stimulated only the Front part of the worm. B) Task 7: Tissue that self-heals after receiving a stimulus. We found a bioelectrical configuration that created a tissue that self-healed after external stimulation. The stimulus started about 5 seconds from the initiation of the experiment, and the V_mem_ of the Front part of the tissue reacted to the stimulus. When the stimulus stopped at about 17 seconds, the V_mem_ of the Front tissue returned to its starting voltage. Other parts of the tissue (Middle and Back) showed no sign of influence from the stimulation. C) Task 8: Tissue that retains V_mem_ (shows memory) after stimulation. We found a bioelectrical configuration that created a tissue that had a memory effect. The tissue received a stimulation about 5 seconds from the initiation of the experiment, and all parts of the tissue reacted to the stimulation. When the stimulus stopped at about 17 seconds, the cells in all parts of the tissue maintained a V_mem_ similar to the one reached during the stimulation. D) and E) are two experiments of Task 9: Finding a tissue that showed a different V_mem_ value for the second stimulus. Both D) and E) stimulated the Front part of the tissue, and all parts of the tissue reacted to the stimulus. D) showed one V_mem_ value for the first stimulation (at about 2 seconds) and a different V_mem_ value for the second stimulus (at about 60 seconds). The Front, Back and Middle regions reacted differently to the stimulations. E) was the same experiment as D), but with different stimulation time and duration of the experiment, to show resilience to different stimulation conditions.

### Task 8: Tissue that retains V_mem_ (memory) after stimulation

The complementary task to a self-healing pattern is one that works as a re-writable memory: once the voltage is changed, can a circuit hold the new pattern? This could potentially be useful not only for synthetic constructs with memory properties, but also to explain phenomena such as two-headed planaria resulting from V_mem_ modulation, which continue to generate two-headed offspring in perpetuity (because their axial polarity pattern memory is permanently re-written to a different pattern by transient stimulation).^52, 53^

We explored the parameter space for a set of parameters that enabled tissue to have long-term memory of a change induced by stimulation. The fitness function gave a higher score to a tissue that stabilized to a resting voltage level, reacted to stimulation, and maintained its new V_mem_ state for the duration of the simulation even after the stimulation had ended,. Because of the relevance to work on planarian regeneration, we used a planarian tissue shape for this task. We found a circuit that started by stabilizing on its resting voltage; then when the external stimulation began, all the cells in the tissue reacted. When the stimulation period was over, all the cells in the tissue maintained their membrane voltage for the rest of the simulation (Figure 9 C). This example used the parameters from the configuration files in Supplement 1 (directory Task 8). We conclude that bioelectric circuit models can explain the re-writability of stable V_mem_ patterns by transient stimuli.

### Task 9: Temporal memory – a tissue that responds differently to first and second stimuli

Finally, we were interested in the ability of bioelectric circuits to perform simple computations and exhibit a kind of temporal memory, by reacting differently to a second stimulation than it reacted to a first instance of stimulation. Numerous examples in biology show this, from preconditioning^54, 55^ to history-dependent regeneration responses in axolotl limbs.^56^ We performed a heuristic search using a fitness function that gave a high score to a tissue that stabilized to a resting voltage and reacted to stimulations. The highest score would be achieved by a tissue in which the average cell V_mem_ during the first round of stimulation was different from that which occurred during the second round of stimulation.

We found a tissue that started by stabilizing on its resting voltage; then when the external stimulation began in the Front part of the tissue, all cells in the tissue reacted. At the end of the first stimulation all cells of the tissue decayed back to the resting potential. When the second stimulation began, the voltage of all the cells reacted to the second stimulation differently. Each tissue part attained a different average voltage from that observed during the first stimulation (Figure 9 D,E). This example used the parameters from the configuration files in Supplement 1 (directory Task 9). These data showed that the cells maintained a sort of memory that caused to the cell to react differently to the second stimulus, maintained in the form of numerical values and the dynamic of the diffusion reaction of the cells. Since the cells did not return to the same conditions they had at the start of the experiment, the history of the activity is still present in the cell dynamics and in the interaction between cells. Thus, the cells exhibited a typical dynamic process that cannot be reversed; so any point in the process is a unique point that contains some clues to the history of the system. We conclude that spatial bioelectric circuits can exhibit memory that distinguishes first and second instances of a specific input.

## Discussion

Bioelectric patterns in tissue are critical determinants of growth and form.^6, 10, 25^ For this reason, efforts to understand the evolution of bodyplans,^57^ to repair anatomy in regenerative medicine contexts,^58^ and to engineer novel biorobotic constructs^39, 58, 59^ will all require control of emergent voltage potential profiles. Biorealistic simulators are now coming on-line for the quantitative modeling of circuit behaviors in silico,^27^ which can be part of a workflow^60^ that solves the inverse problem: what set of ion channels and gap junctions, if expressed in cells, lead to the emergence of specific bioelectric prepatterns? Here, we report an initial effort toward applying machine learning tools to the discovery of initial conditions (ion channel parameters) that form a circuit with desired spatio-temporal properties.

We focused largely on spatial patterns, because developmental bioelectric signaling requires slowly changing geometric prepatterns. However, many examples of more neural-like behavior (spontaneous firing, propagating waves, etc.) were also observed and can be modeled using this method. One major knowledge gap that we sought to address was exploration of the space of possible bioelectric circuits. As published work in the field characterizes endogenous bioelectric circuits and their resulting patterns, it is important (for evolutionary and synthetic bioengineering purposes) to also understand the latent space around these specific parameters. What can bioelectric circuits do – what general behaviors can possibly be found, and what search methods are promising approaches for identifying them?

We chose genetic algorithms for heuristic search, among other reasons, because these algorithms approximate the natural process that led to the emergence of multicellular anatomies. We searched the space of ion channel parameters, which in biology are set by the genes encoding each ion channel and gap junction protein. The dynamics we observe shed light on the kinds of dynamics that are achievable by an evolutionary process operating over the space of ion channel-encoding genes and the resulting multicellular voltage patterns.

### Limitations

From our experience with the tasks above and from prior work,^61, 62^ there are at least three major obstacles to the effort of identifying desired bioelectric circuit parameters. First, heuristic searches in general, and genetic algorithms in particular, assume that the parameter space can be understood and interpreted as gradually changing toward the best possible solution. The bioelectric phenotype space has not yet been mapped in detail, and it was unclear how rugged or amenable to hill-climbing search it would be. Second, the heuristic search also assumes that the fitness function can guide the search to the target by following the highest fitness score. If the space of the fitness score is a nonuniform distribution of scores, then the heuristic search can drive the search in unexpected directions. Furthermore, if the fitness space does not have a clear gradient, the expectation is that the fitness function should know to handle the distorted fitness score space and provide a gradual path toward the target, regardless of the terrain. Third, defining the desired target is also a challenge. For example, in task 5, we wanted to search for a tissue that has the same V_mem_ features as found in the bullseye image. We designed a fitness function that gives a score to a tissue that shows a significant gradient from the center of the tissue to the outer border of the tissue. This experiment did not find precisely what we were looking for; however, one of the patterns found (see Supplementary images) shows an image similar to one of the possible images of a bullseye. That reveals that similar to the alignment problem in machine learning ^63^, designing a fitness function even for a simple pattern like a bullseye can be a non-trivial task, because many other possibilities may resemble the desired target, and it’s difficult to formalize a function that captures all the features that a human observer intuitively recognizes as correct (i.e., shifted, scaled, and slightly distorted versions of the ideal pattern).

### Next steps: the future of bioelectric circuit design and discovery

Our findings suggest that the bioelectric parameter space contains many different patterns, where each one of the patterns resides within different sizes of islands that contain some variation of the same pattern. Some of the islands have the potential to be candidates for a given target and others are not. This is difficult to illustrate, because the search involves 33 dimensions, while other studies that use a simpler model like a Hodgkin-Huxley neuron use fewer parameters (e.g., like those found in ^62, 64^, which show how a typical parameter space can be visualized). We also explored only a small proportion of the parameter space, since 33 dimensions constitute quite a large space to experiment and can easily rival the number of atoms in the universe. Each configuration file has 33 different parameters that the heuristic search can manipulate. For simplicity, let’s assume that each parameter has only 100 possible values; that would give us 100^33^ = 1.e+66 possible parameters to explore. Running BETSE takes about 50 minutes on a strong computer, which makes for a very time-consuming task overall. We expect that with improvements in high-performance computing, much more high-resolution searches can become tractable.

There are several different paths to explore for more effective searches in the future. Using the result of our current exploration, we hope to devise a machine learning method that can speed up the heuristic search by leveraging the experience of past explorations to improve the gene selection process and the fitness function. Machine learning can steer the search to areas where there is more chance to find the desired pattern. Another option is to add some intelligence to each cell, maybe using embedded Gene Regulatory Network elements inside each cell to shift part of patterning formation and cell intelligence to the cell and begin to implement a more life-like multi-scale competency architecture.^65, 66^ Finally, it may be possible to add a translation layer between the parameters exposed to the heuristic search and the cell parameters (ion channels, conductance level, etc.) – in effect, enriching the physiological information-processing. This additional layer will translate the heuristic search parameter tuning to a non-polynomial change of the underline cell parameters. A change in one of the heuristic search parameters could transform this translation layer into a series of ion channel manipulations in a non-polynomial way. Thus, the translation layer will try to correct the deformities in the rugged fitness space, by shifting part of the work to the cell and its interaction between other cells. A final future direction is to design a fitness function that interacts with the environment, similar to the interaction between actor-critic artificial neural networks.^67^

Regardless of the details, it is very likely that the incredibly rapid pace of advances in parallel computing and machine learning will impact this field positively over the near future. We envision sophisticated artificial intelligence systems that will work together with human scientists to enable a new generation of bioinformatics, complementing the current focus on biochemical and transcriptional circuits with the extremely powerful machinery of bioelectrics. This will have enormous implications for understanding and rational control of voltage-based signaling in health, disease, and synthetic bioengineering applications.

## Acknowledgements

We thank Julia Poirier for assistance with the manuscript, and Alexis Pietak for assistance with the use of BETSE and related aspects of the research. We acknowledge the use of the Tufts University High Performance Compute Cluster (https://it.tufts.edu/high-performance-computing) and the assistance of the HPC team. This work was supported by a sponsored research agreement from Morphoceuticals Inc.

## Supplemental Figures

**Supplement Figure 1.**
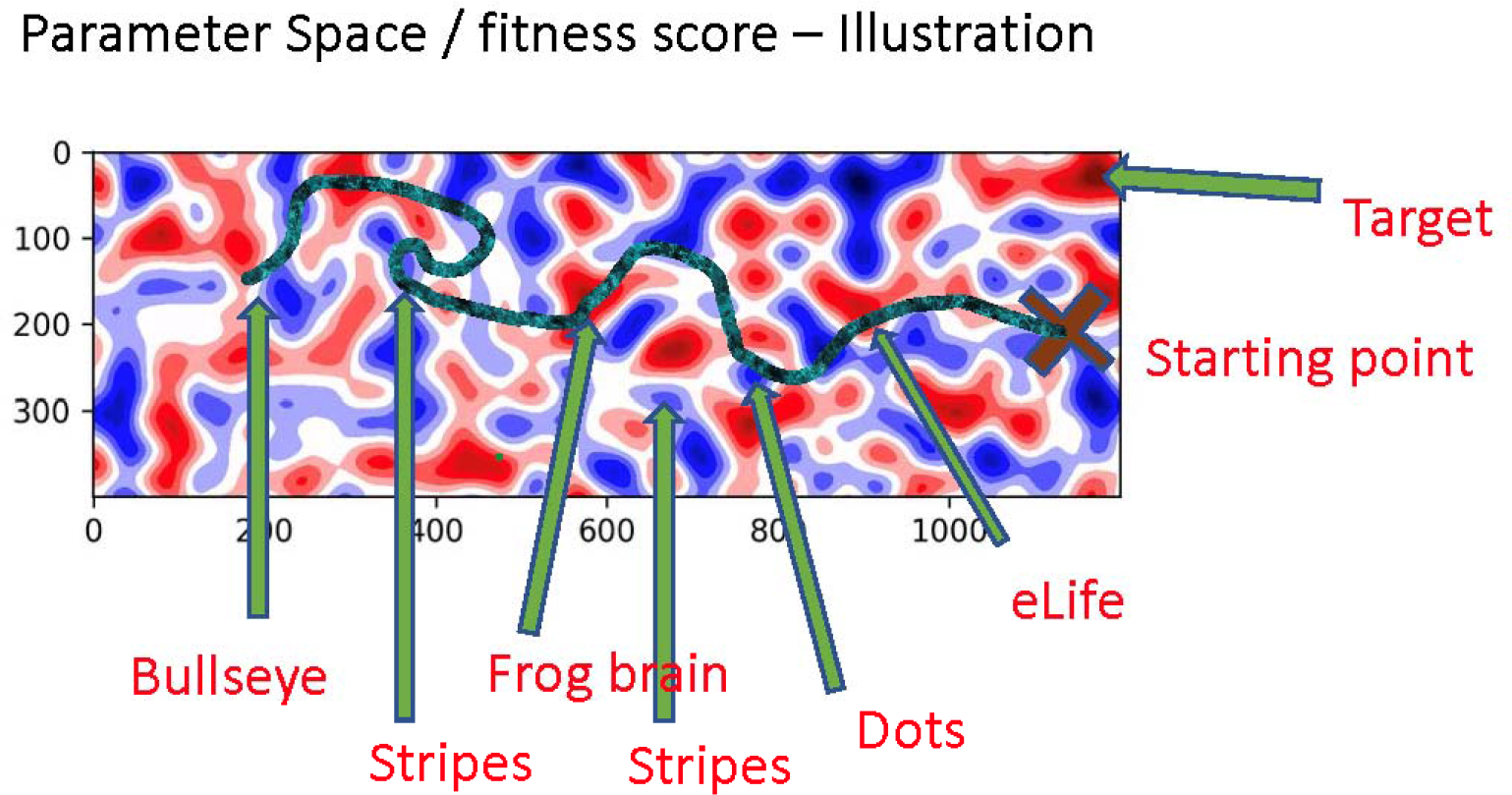

**Supplement Figure 2.**
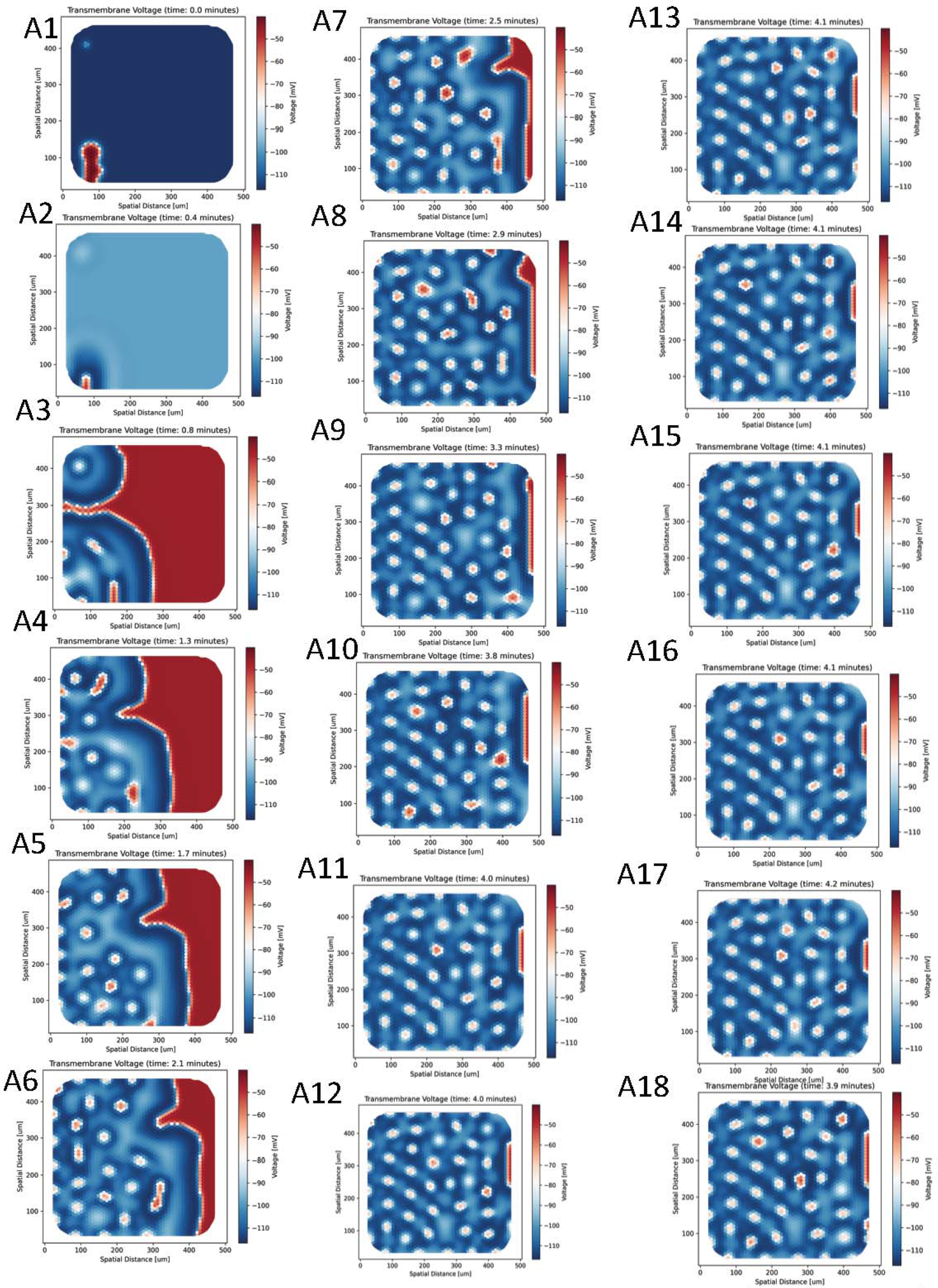

**Supplement Figure 3.**
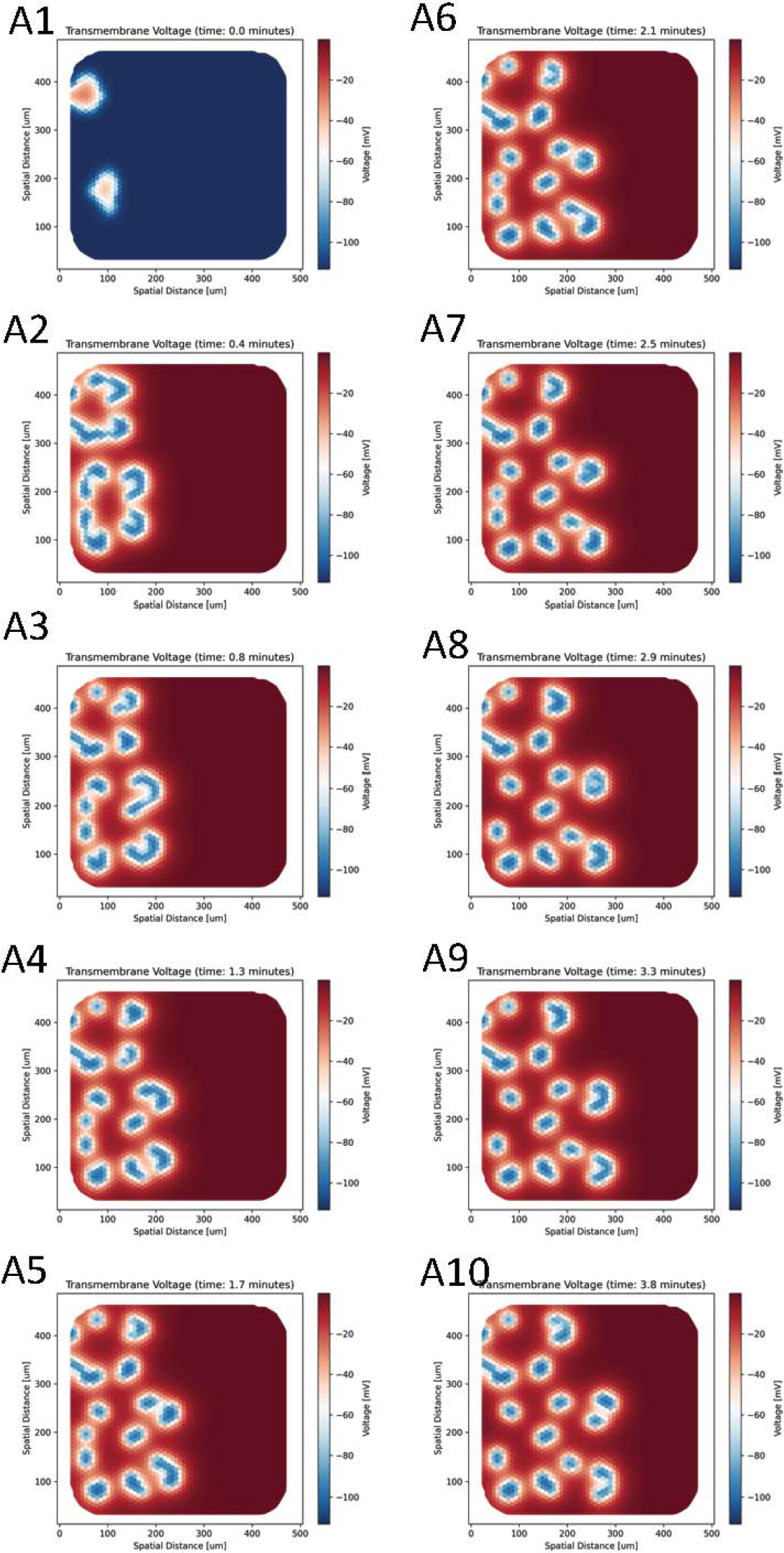

**Supplement Figure 4.**
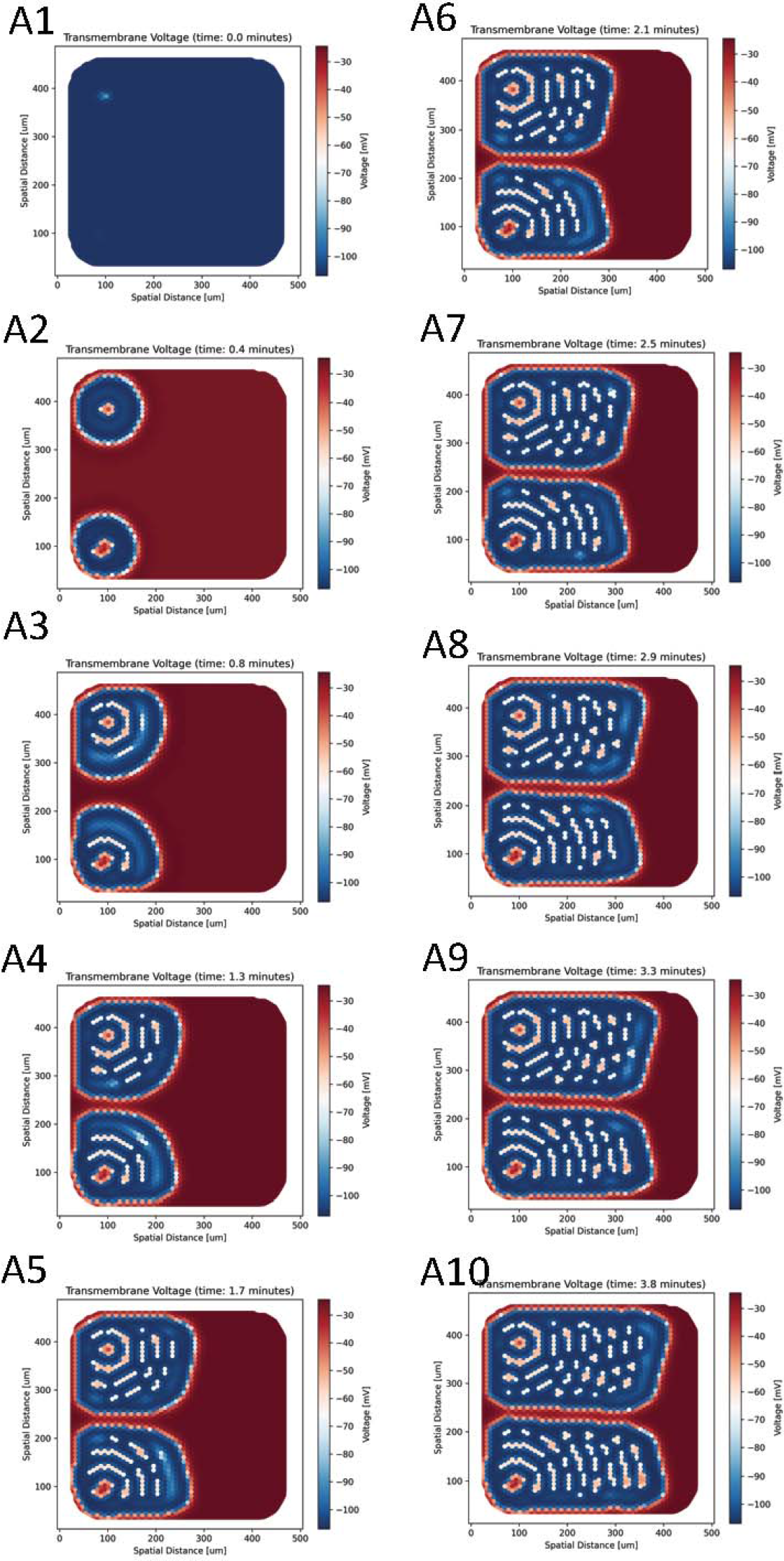

**Supplement Figure 5.**
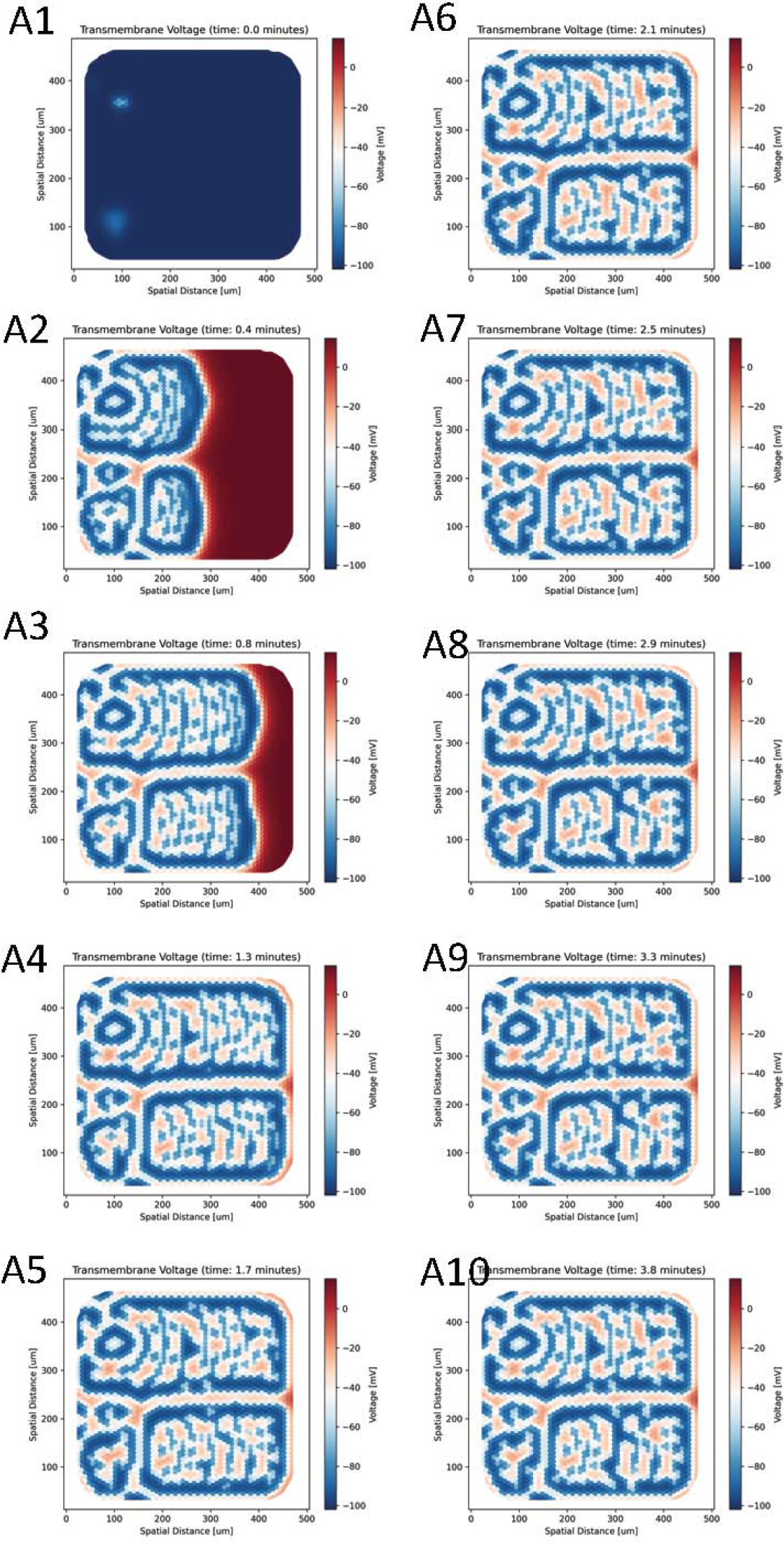

**Supplement Figure 6.**
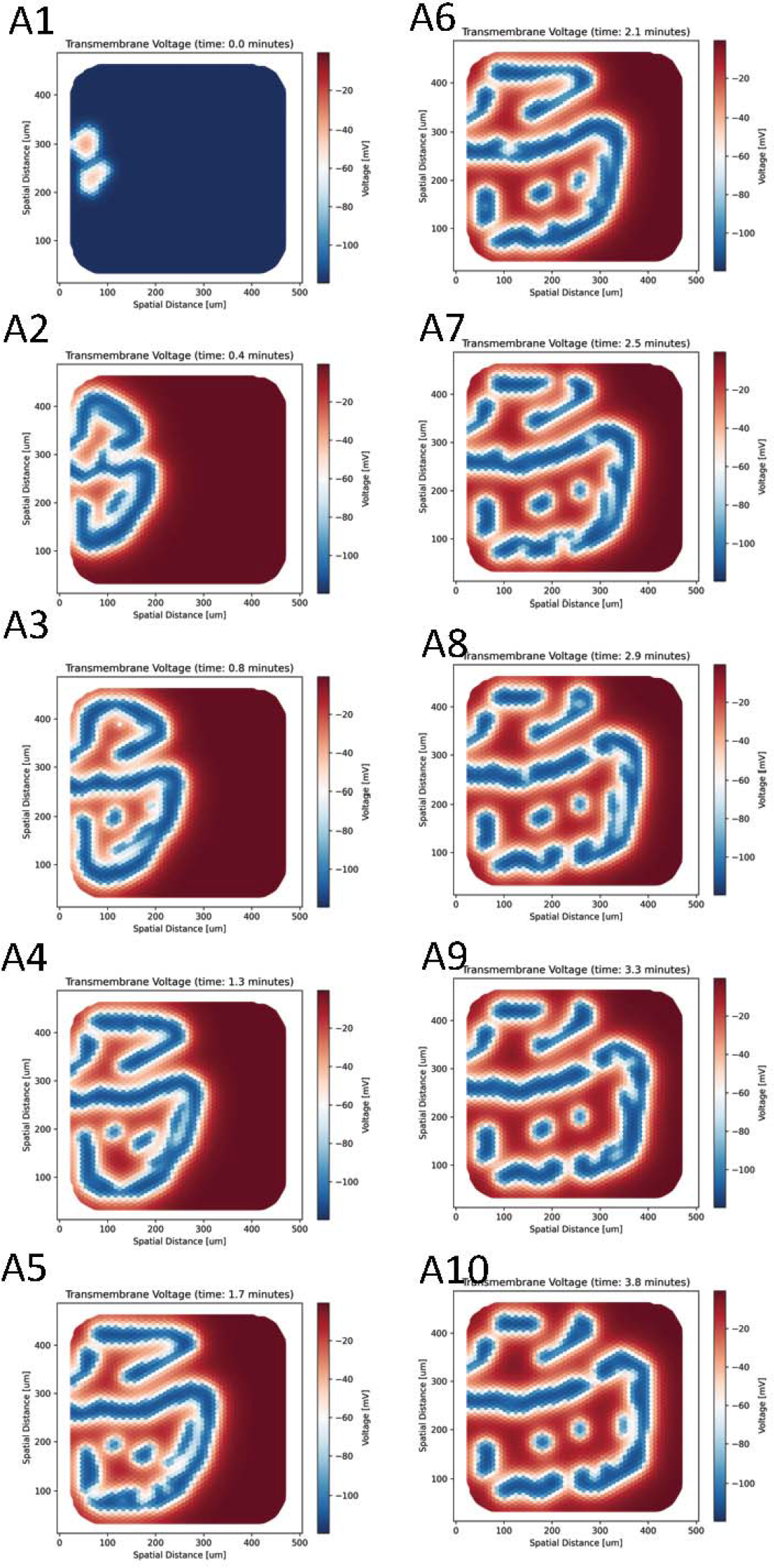

**Supplement Figure 7.**
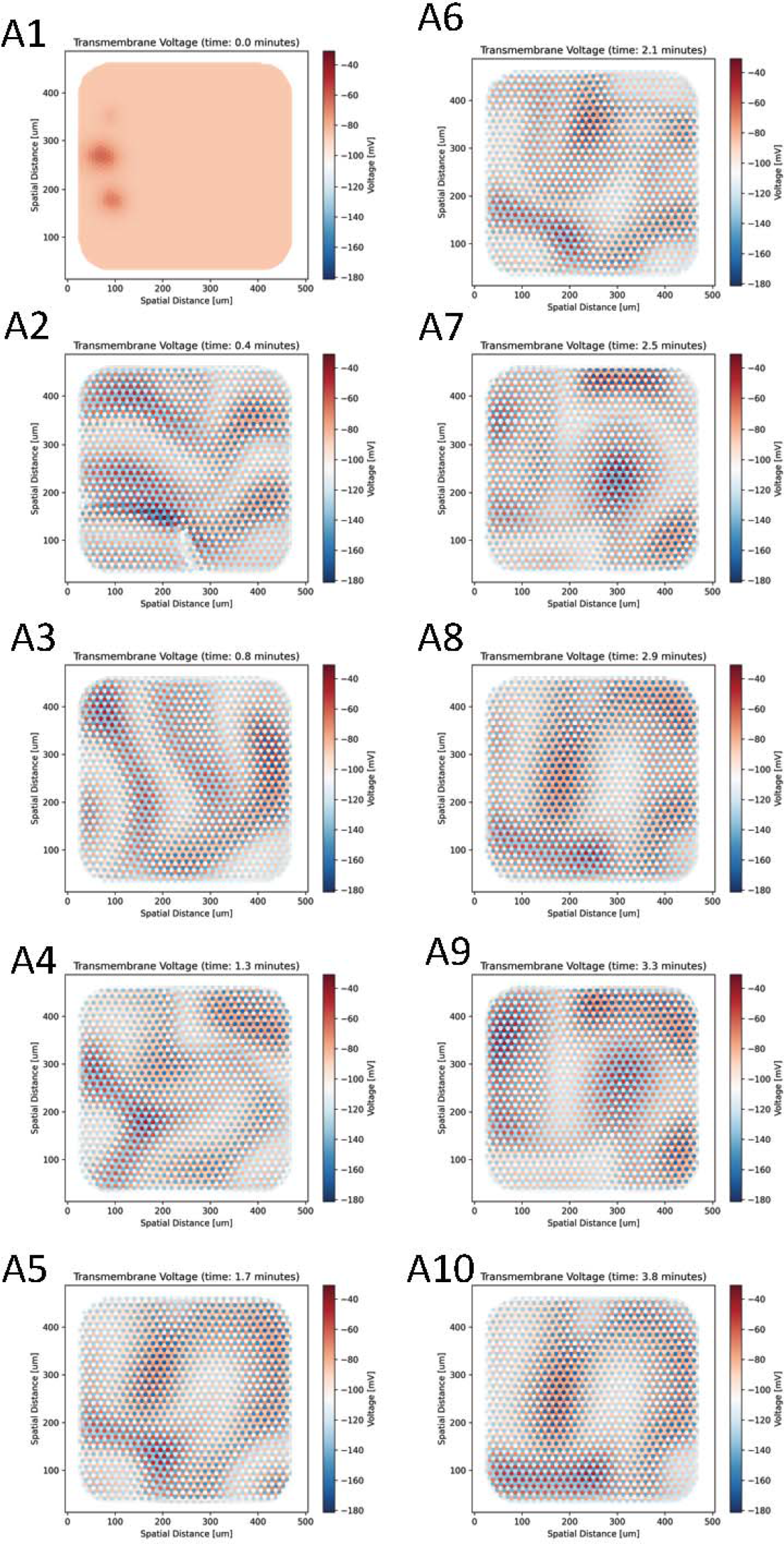

**Supplement Figure 8.**
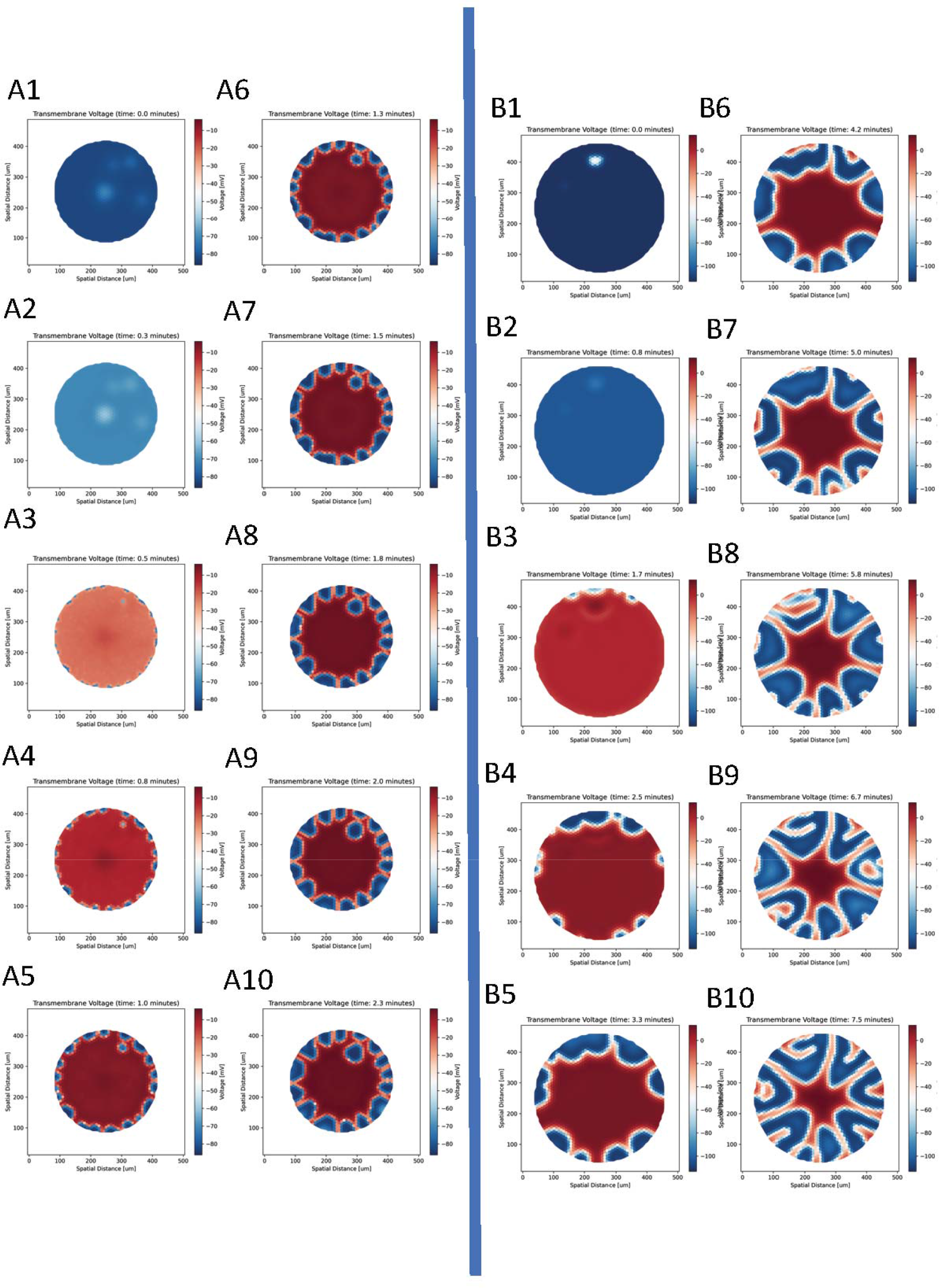

**Supplement Figure 9.**
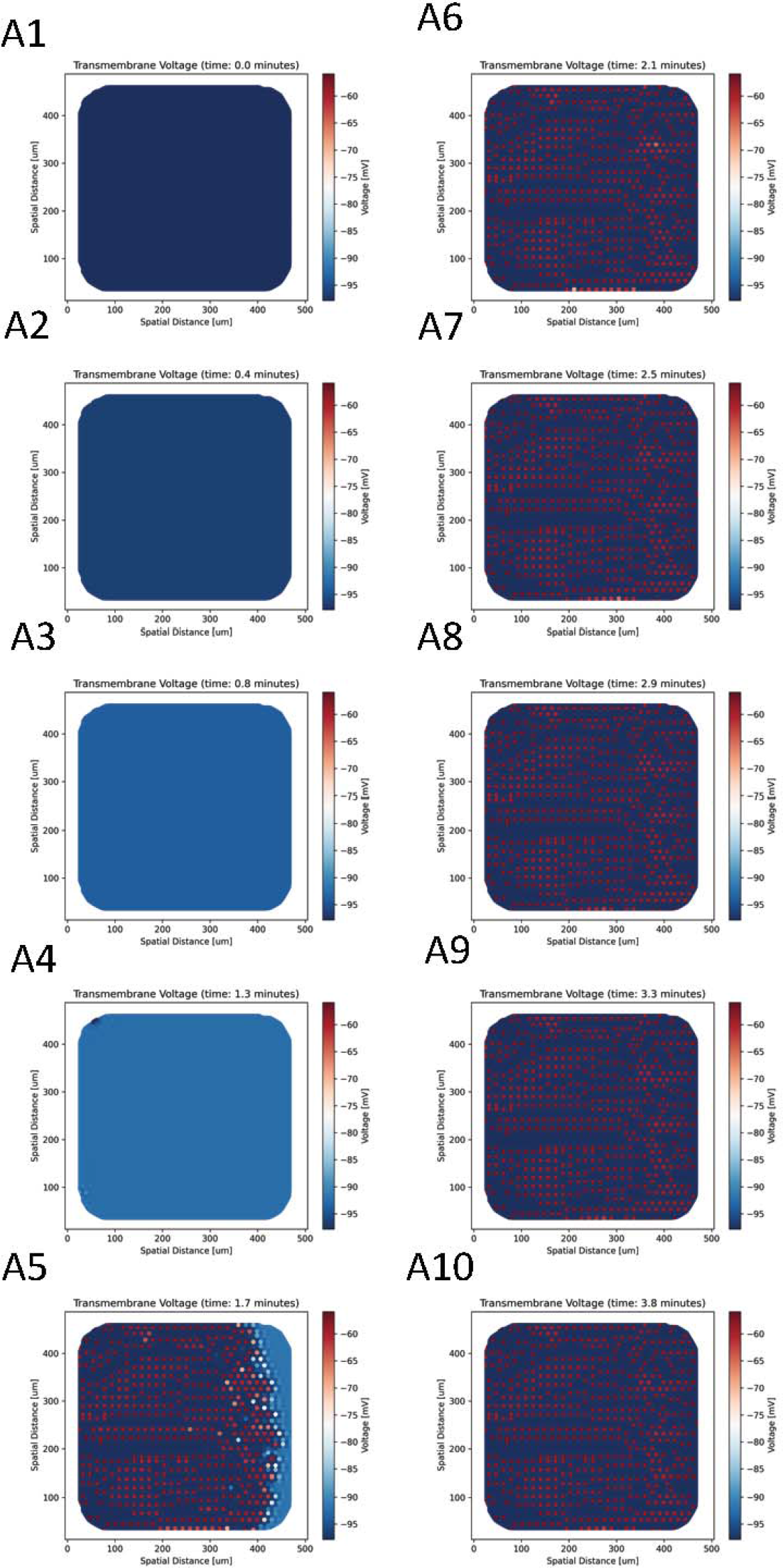

**Supplement Figure 10.**
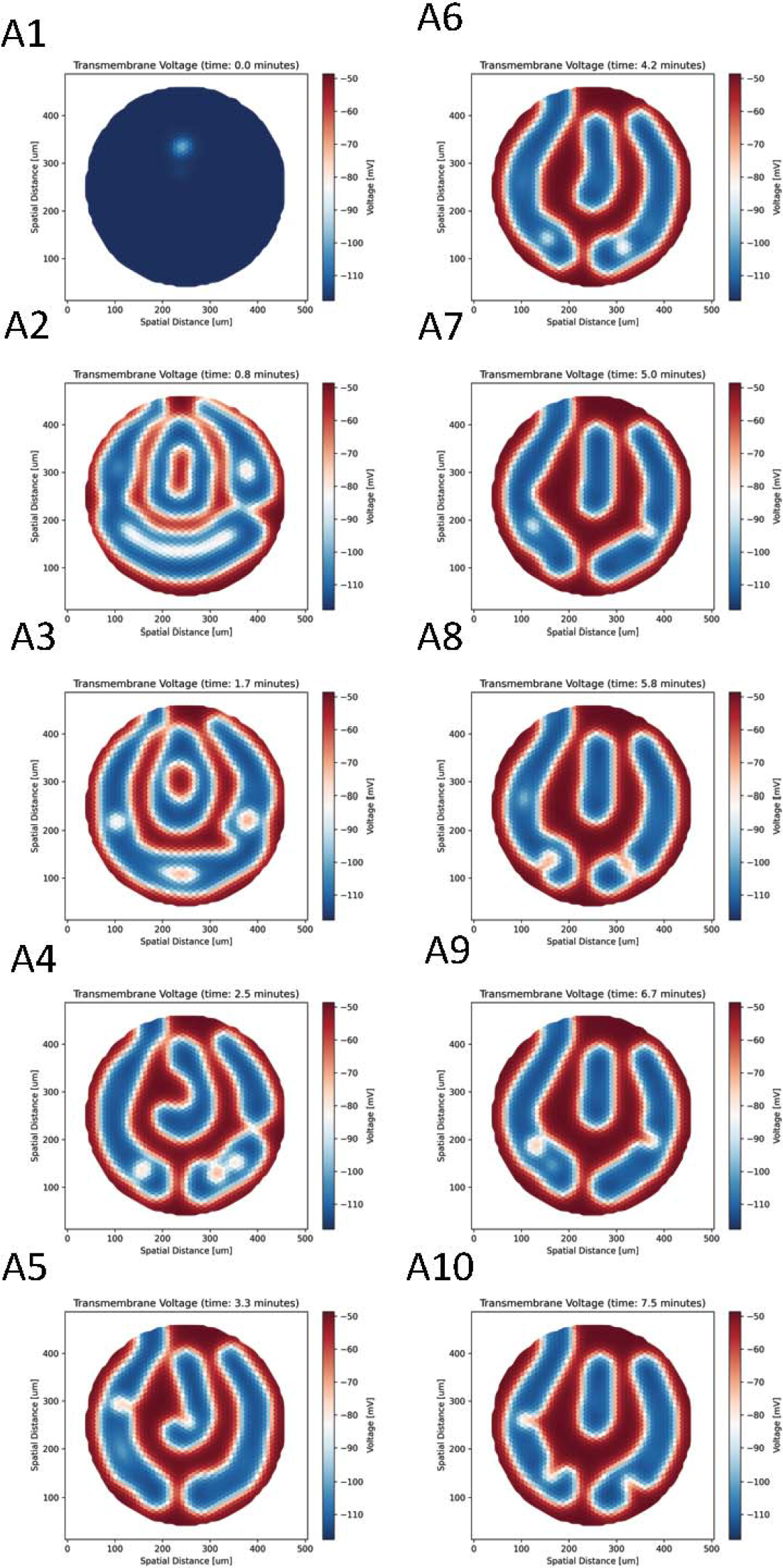

**Supplement Figure 11.**
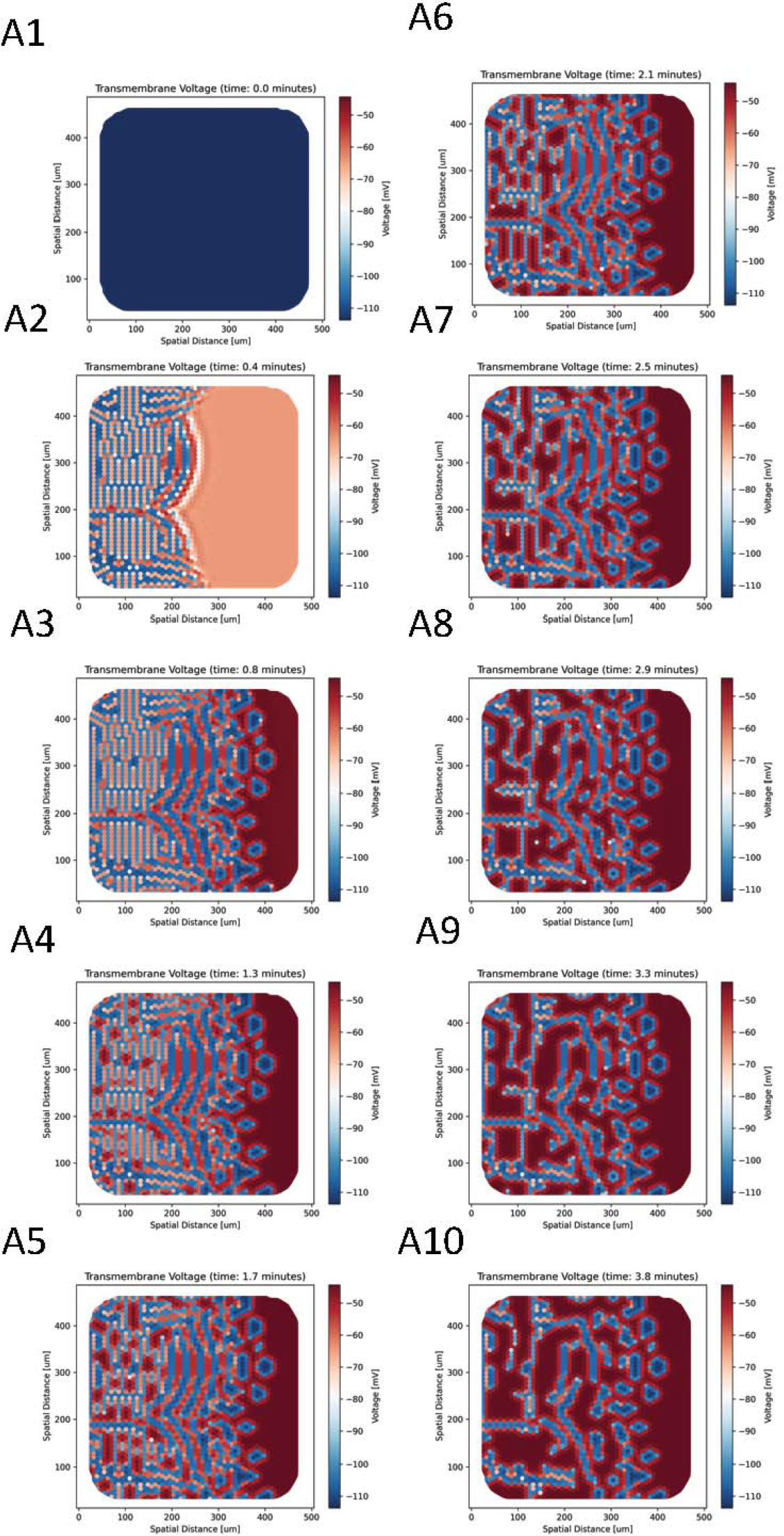

**Supplement Figure 12.**
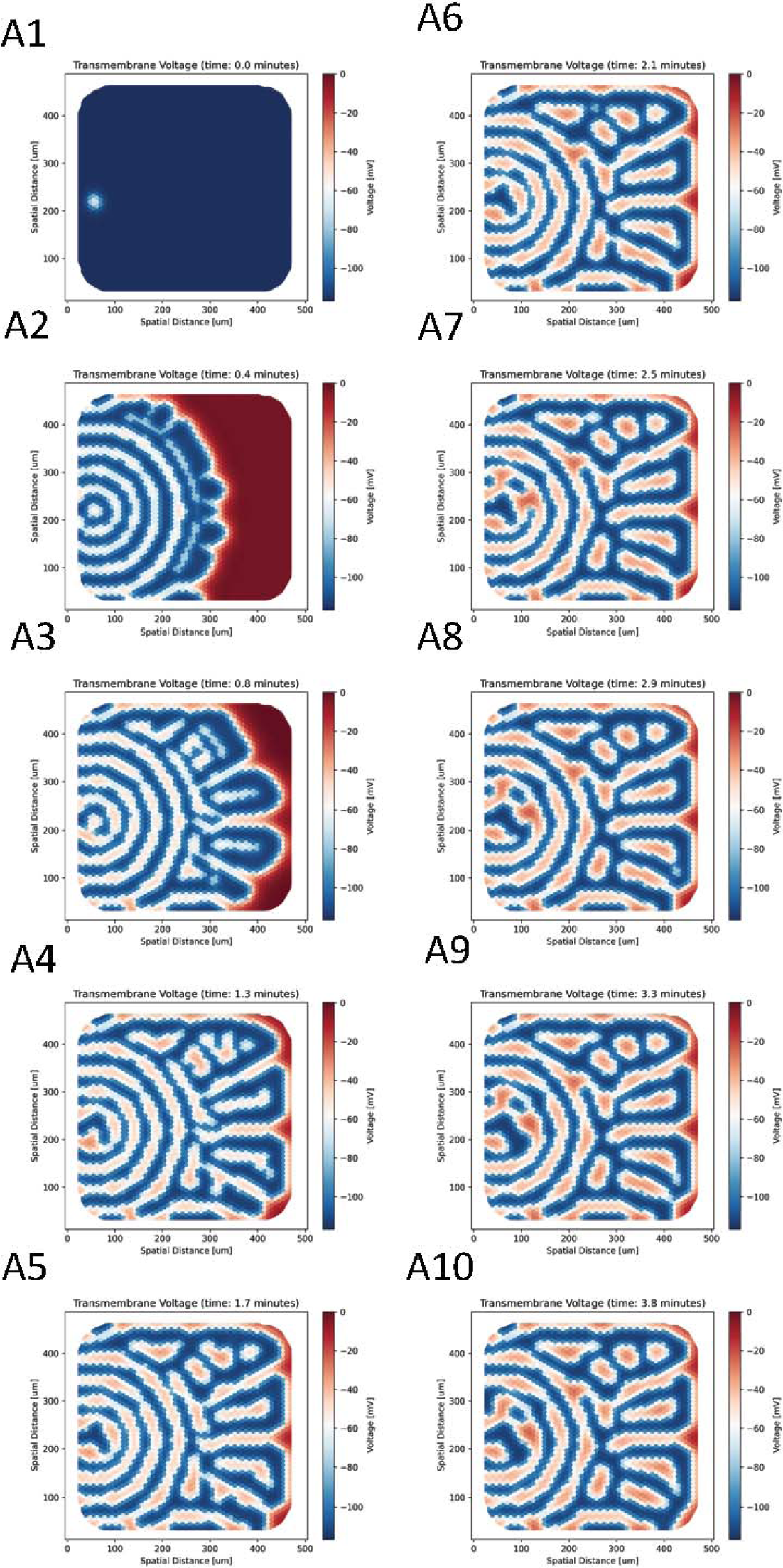

**Supplement Figure 13.**
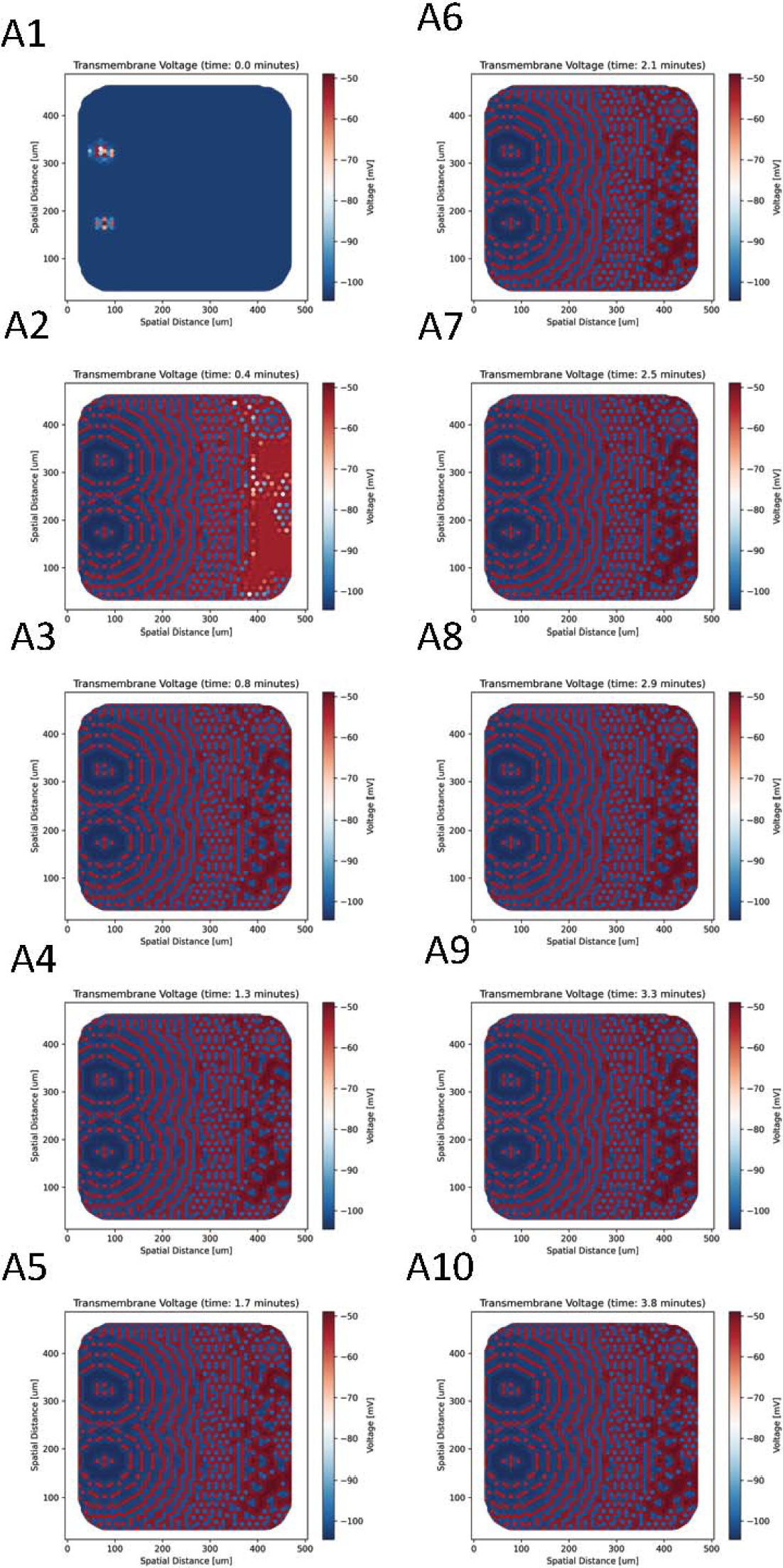

**Supplement Figure 14.**
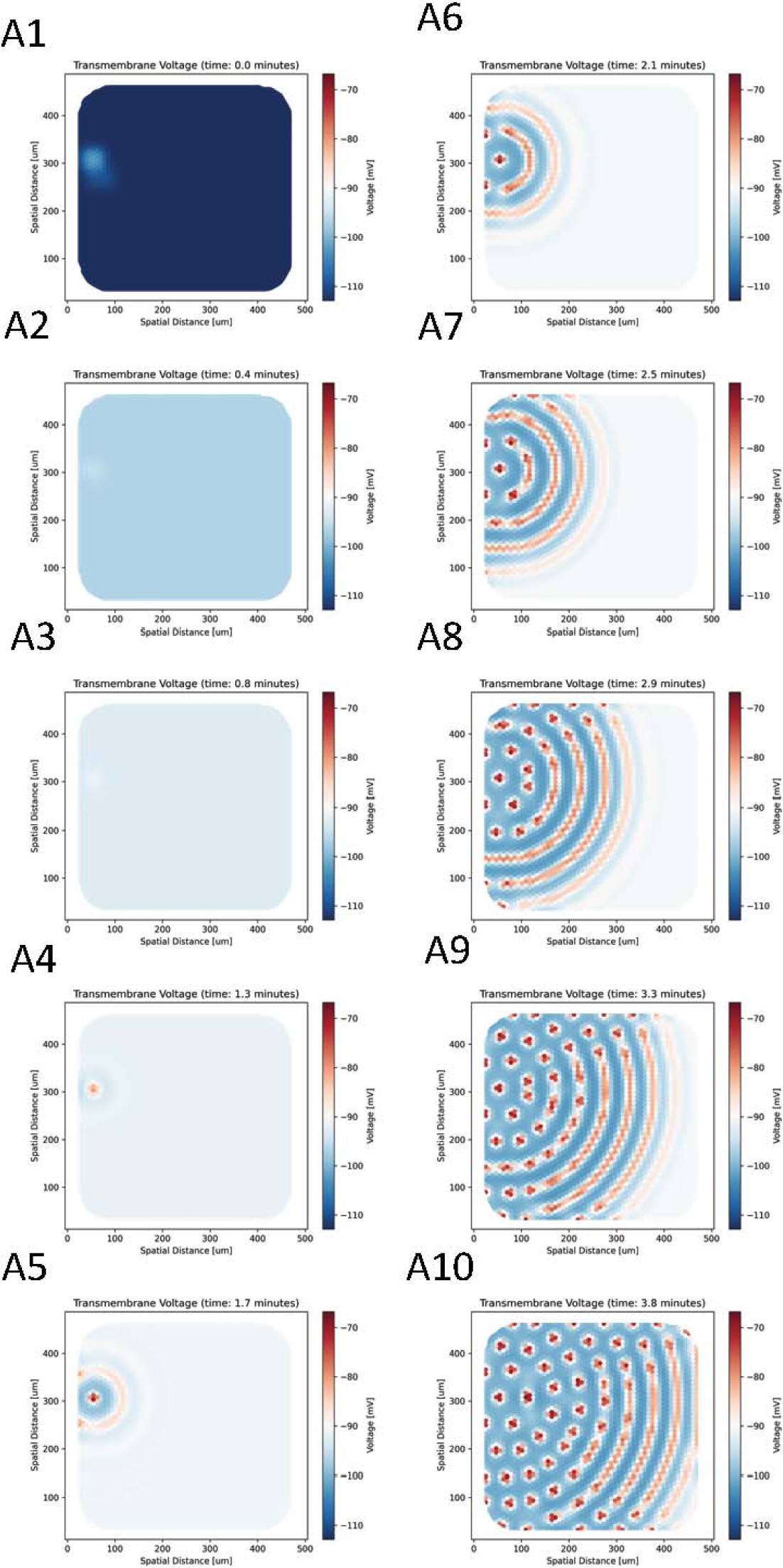

**Supplement Figure 15.**
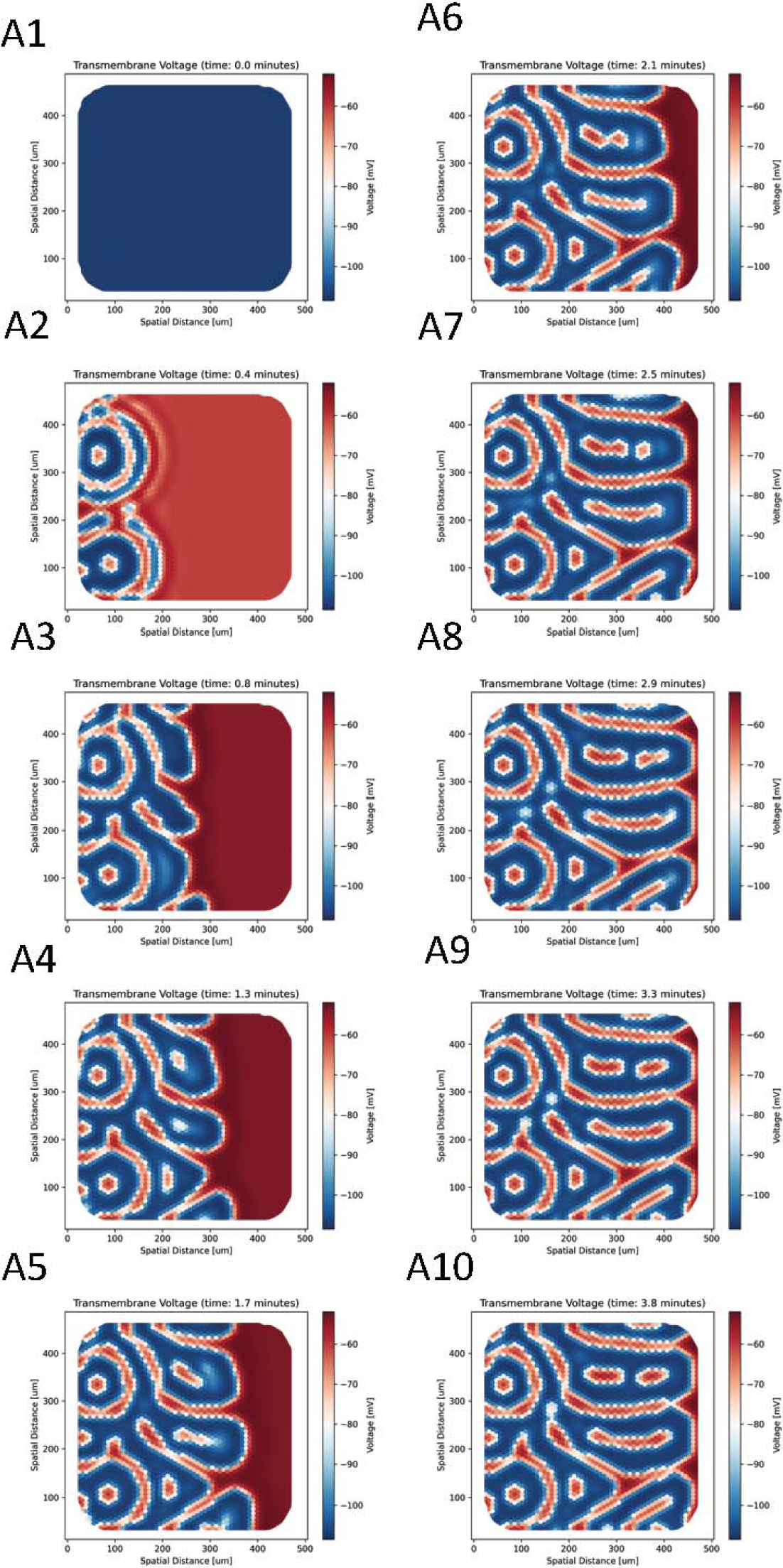

**Supplement Figure 16.**
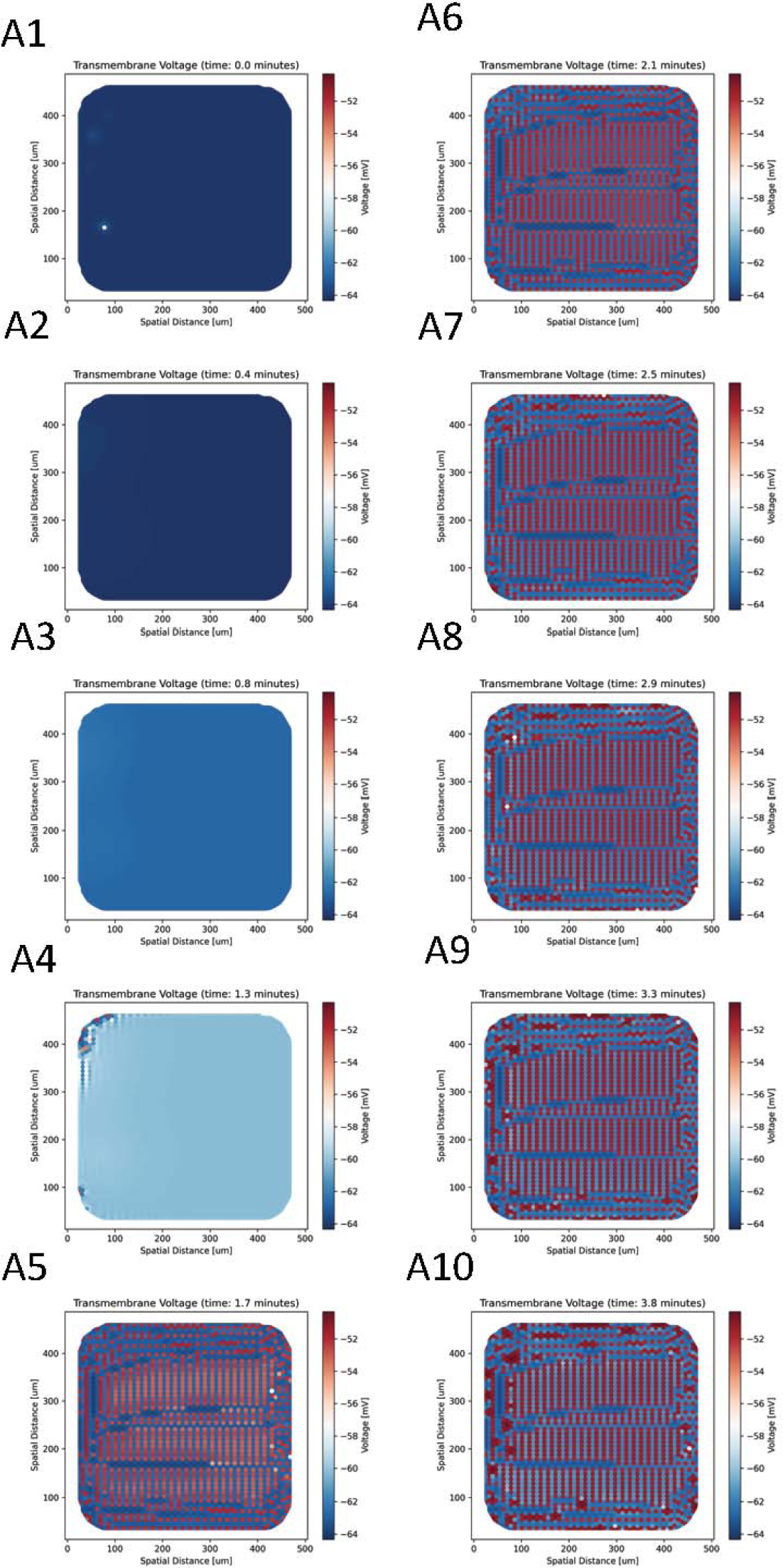

**Supplement Figure 17.**
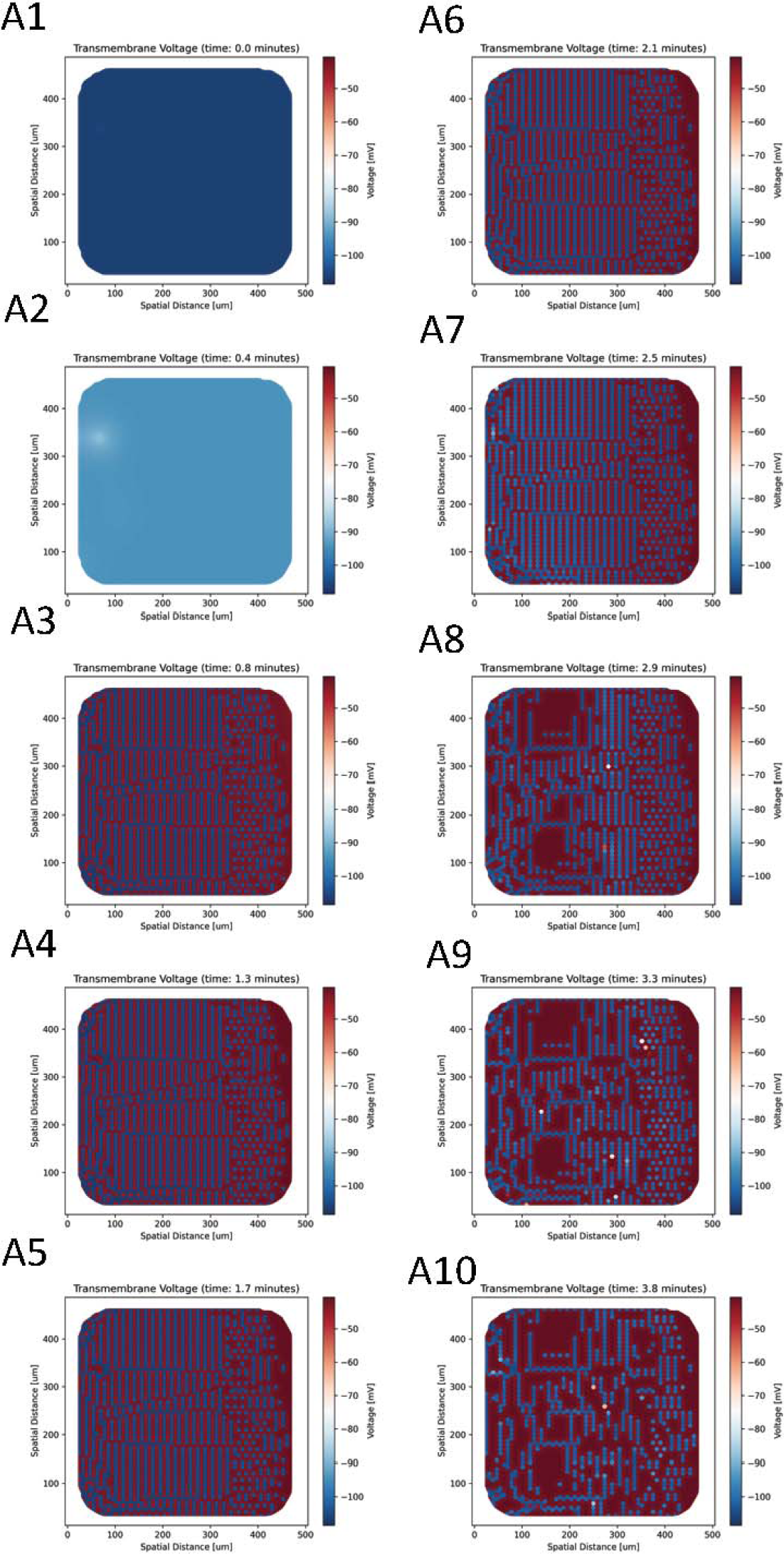

**Supplement Figure 18.**
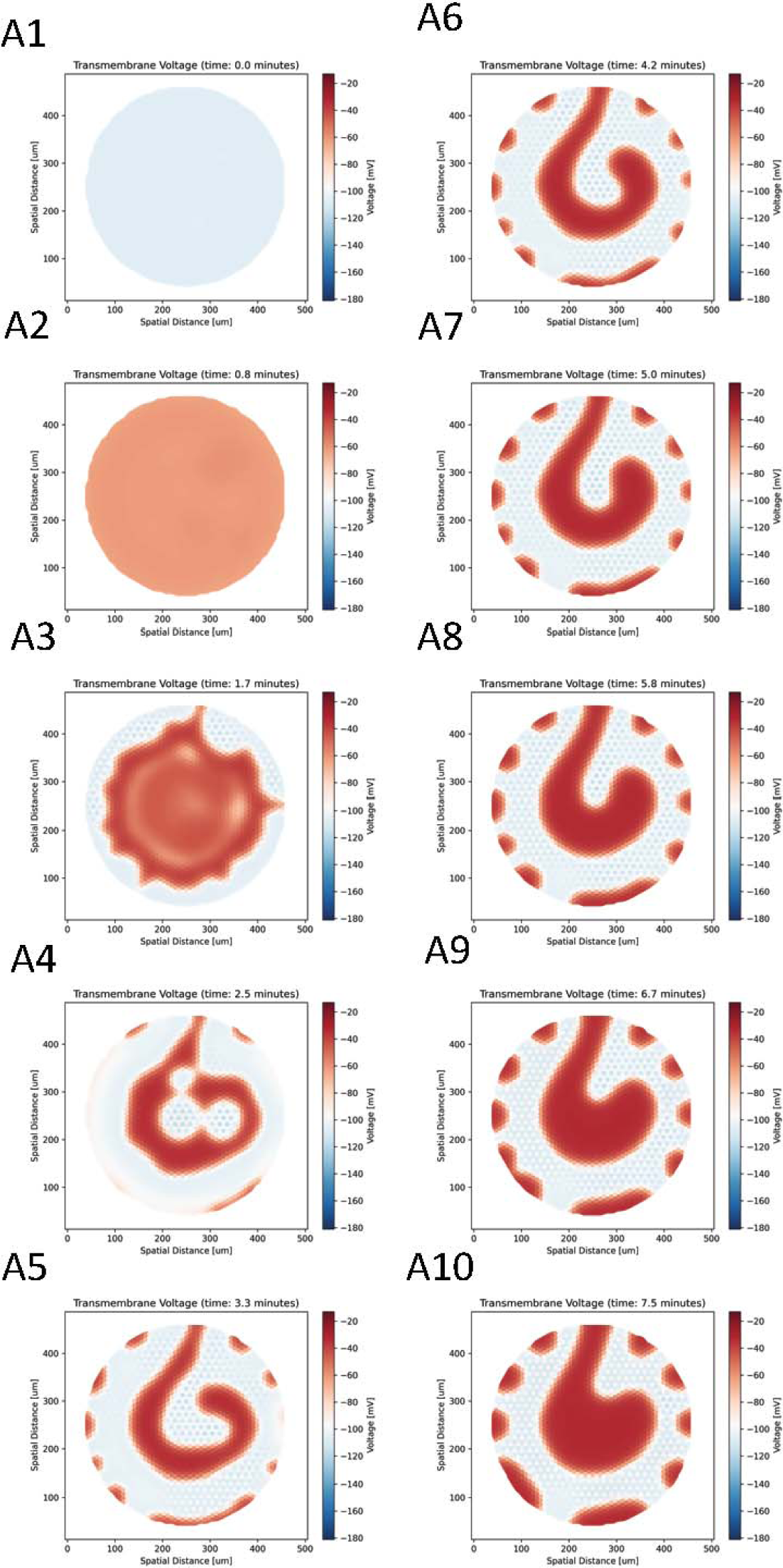

**Supplement Figure 19.**
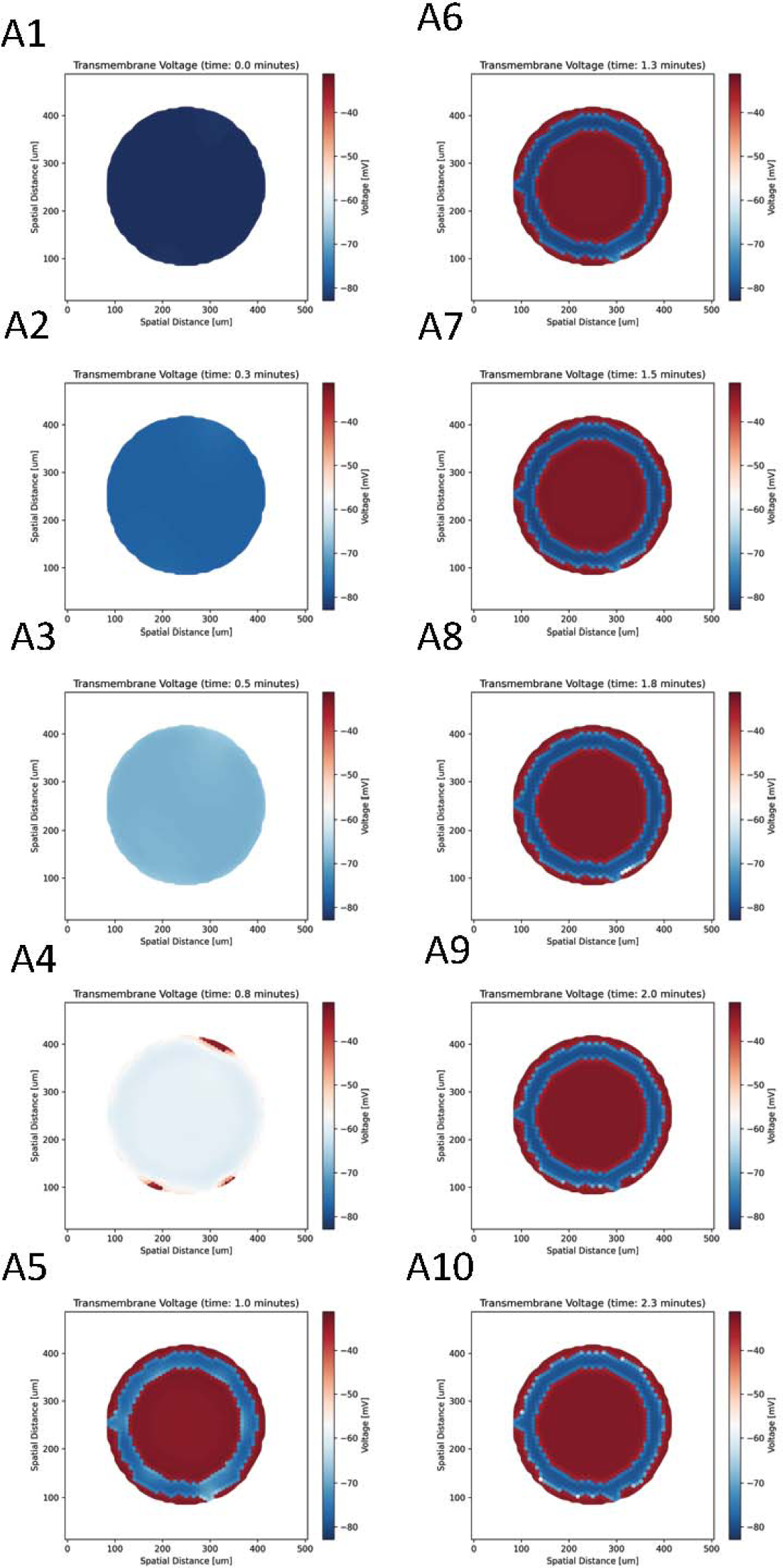

**Supplement Figure 20.**
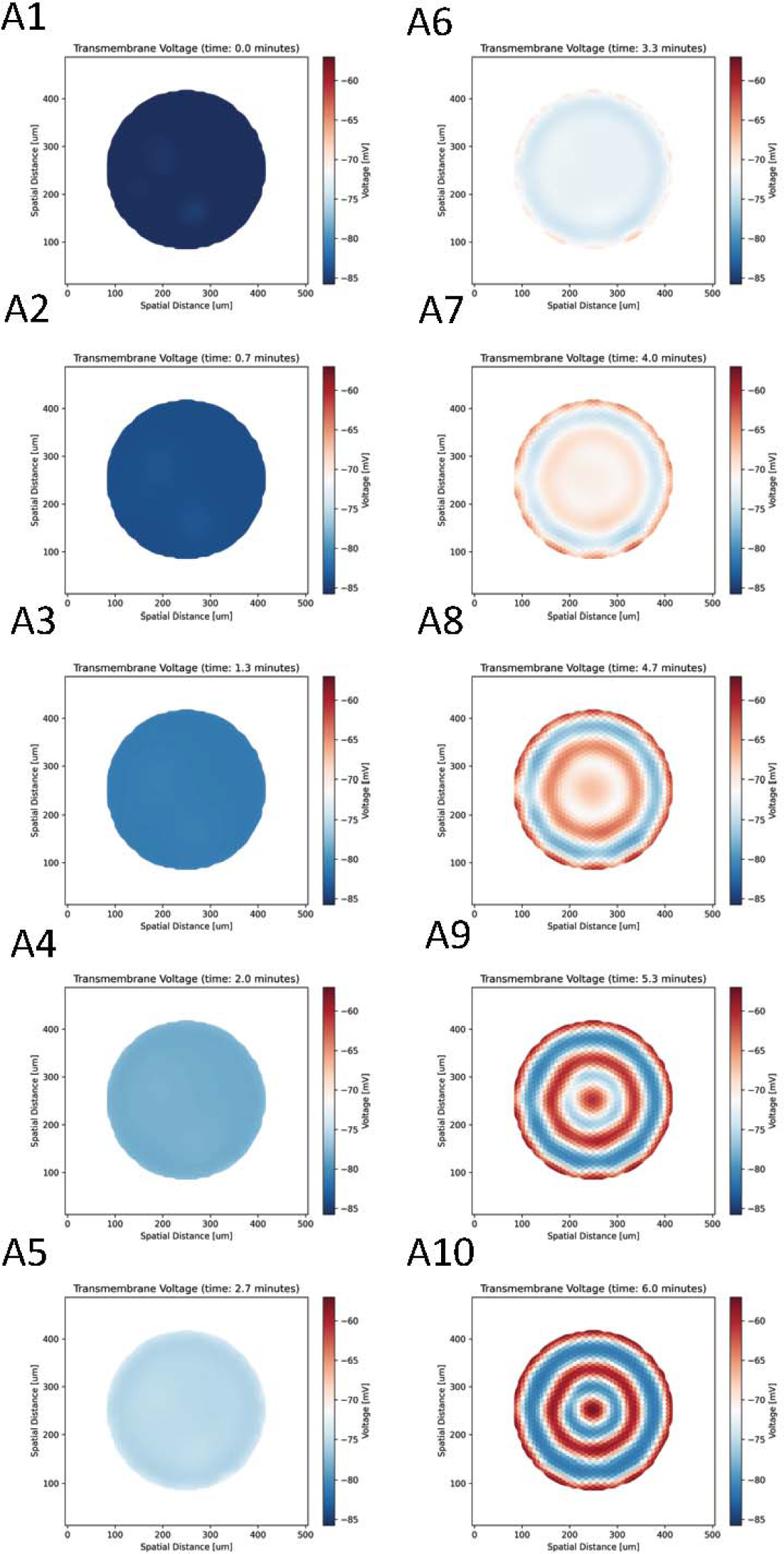

**Supplement Figure 21.**
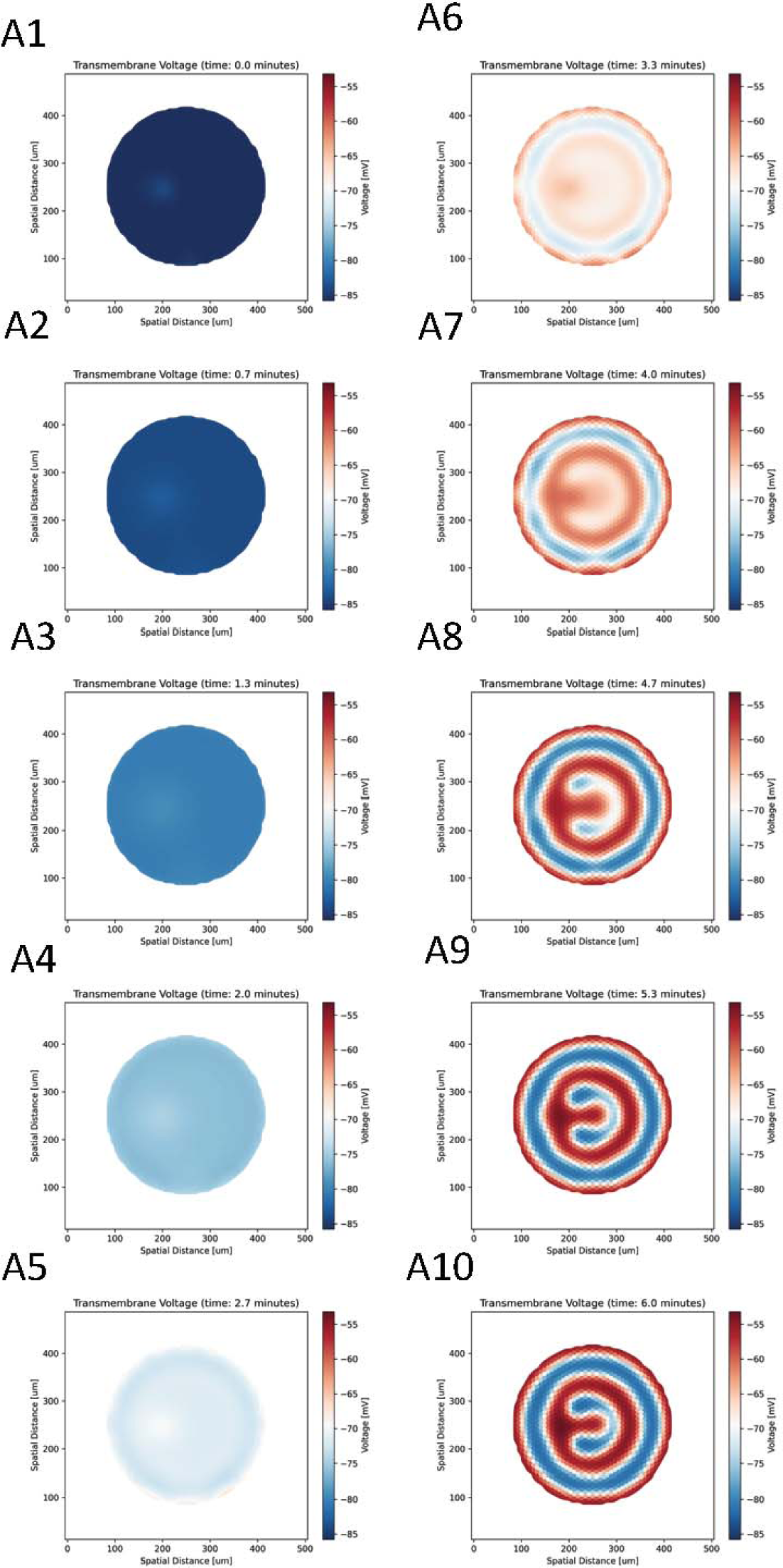

**Supplement Figure 22.**
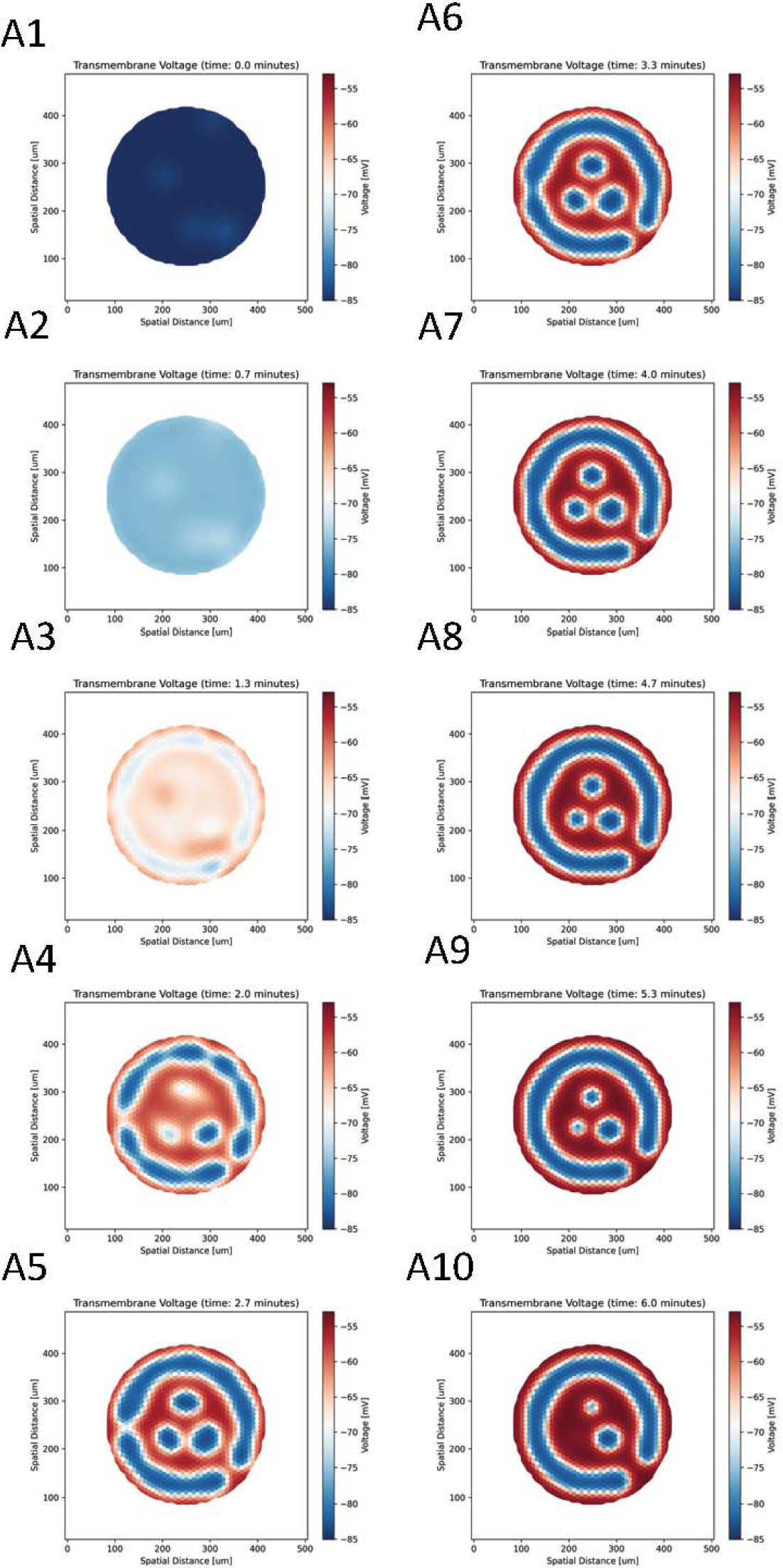

**Supplement Figure 23.**
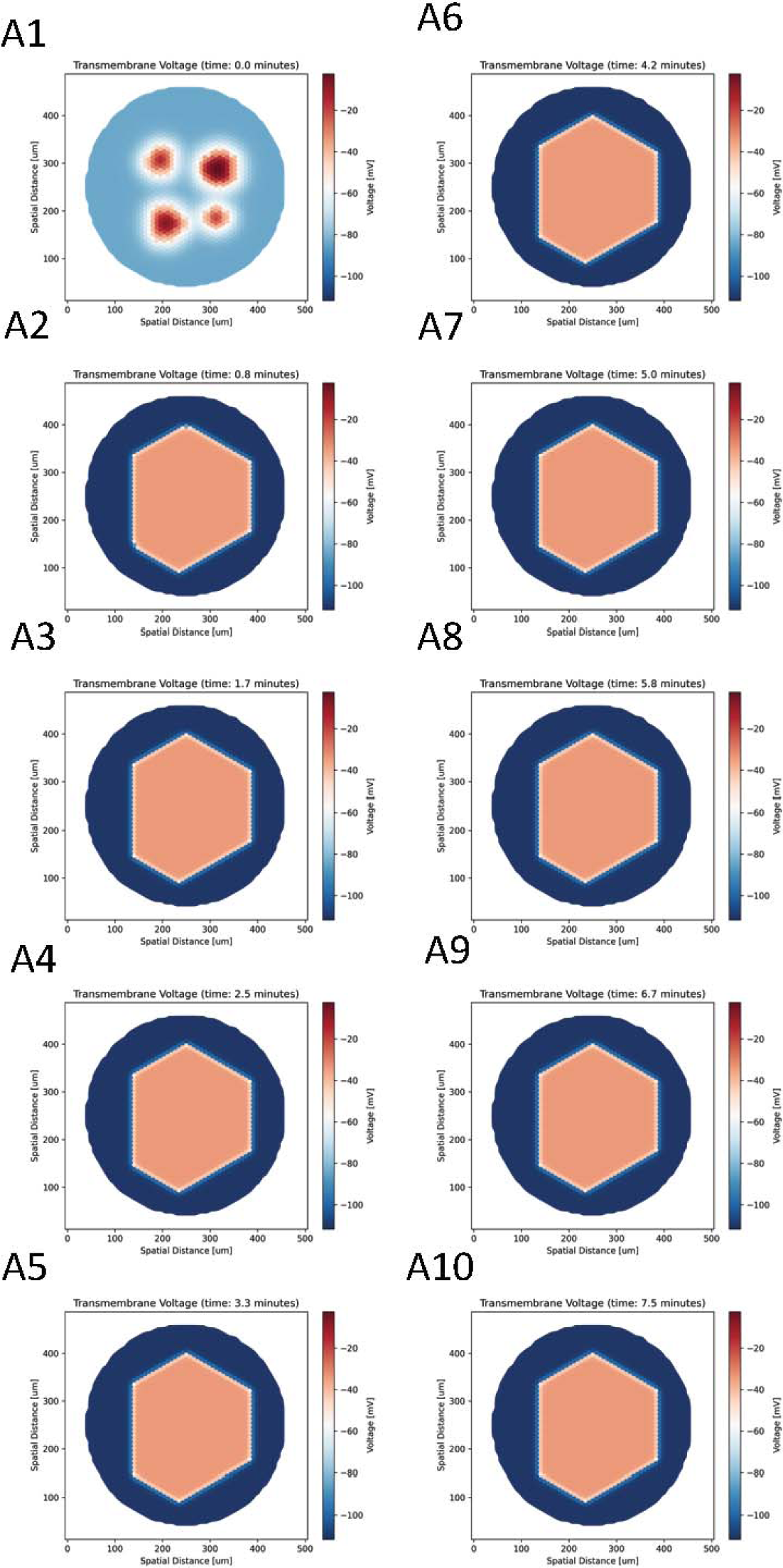

