## Supplemental Materials for "Exploring The Behavior of Bioelectric Circuits using Evolution Heuristic Search"

All supplemental materials for this paper can be retrieved from https://dataverse.harvard.edu/dataset.xhtml?persistentId=doi:10.7910/DVN/U1QRS8

Configuration values for each experiment

In order to recreate the experiments in this paper, we assume the reader has the basic knowledge on how to use the BETSE (https://github.com/betsee/betse) simulator. Our supplementary materials include all the configurations needed to run BETSE to produce each of the bioelectrical patterns in this paper. The file structure of supplementary materials folder (Paper - BioElectric - BETSE – configs) is:

/

├───Task 1 - Vmem change as little as possible over time

├───Task 2 - Tissues that have as little change as possible through time, but with high variance between cells

├───Task 3 - Fit specific Vmem

├───Task 4 - High variance between Vmem cells

├───Task 5 – Smiley and bullseye

├───Task 6 - Patterns with insensitivity to shape or size of tissue

├───Task 7 - Tissue that self-heal when a few cells getting input

├───Task 8 - Tissue that retain Vmem (memory) after stimulation

├───Task 9 - cells that respond similar to flip flop operation turn out to be form of memory using chaos

└───Interesting patterns

│	├───Configurations

│	├───Image Frames

│	└───Movies

| TOC

Tasks 1-9

Each of the Task 1 to Task 9 directories contains configuration files for the corresponding Task (as described in this paper). Within each Task directory there is at least one top-level .yaml file; this is the configuration file that should be fed into BETSE to reproduce that Task. Some Task directories also have additional subfolders (e.g., extra_configs, geo) that will automatically be discovered and used by BETSE, as directed by the aforementioned .yaml configuration file.

Interesting patterns

We had assumed that our parameter/fitness space would have smooth gradual properties that would help our heuristic search algorithm to progress toward the desire morphological target. However, we found that our naïve approach presented a parameter space that was too confusing for exploration by our heuristic algorithm. We did encounter many “interesting patterns”; but rather than occurring along a continually increasing gradient of fitness score, these patterns presented “islands” of fitness, as shown by the peaks and valleys in the illustration below (Supplementary Figure 1).

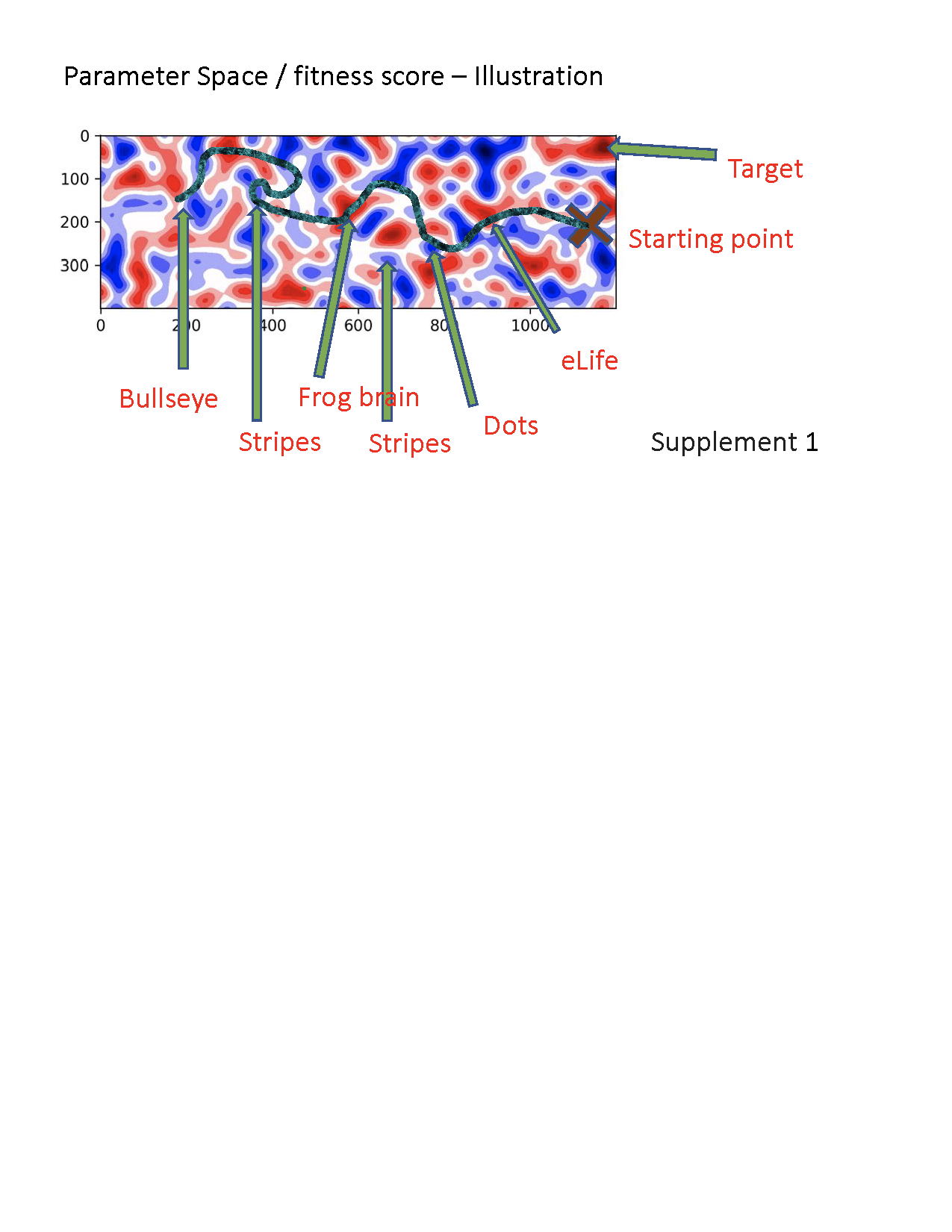

This proved to be problematic, because our heuristic algorithm required a steadily increasing fitness gradient in order to progress smoothly to the target morphology. The presence of morphologically “interesting” islands (peaks or valleys) interrupted what we had hoped would be a smoothly increasing gradient. The result was that the heuristic algorithm could “get stuck” in particular areas along the serpentine line of fitness scores, effectively stopping it from further exploration of other “islands” along that line.

The “Interesting Patterns” directory contains 3 subdirectories – Configuration, Image Frames, and Movies – as well as a Table of Contents file. The “Image Frames” directory contains forms from the tissue process in the experiment. The “Movies” directory contains the full movies for each experiment.

The TOC excel file contains 32 rows, each of which contains an image of the target tissue morphology (the “interesting pattern”), a description of fitness target that was used by the heuristic search, and the serial number of the experiment that produced that pattern. This serial number corresponds to the name of a subdirectory under Configurations.

If the reader wants to reproduce the image frames and the movies for any particular “interesting pattern” shown in the TOC, s/he should go to the appropriately named Configurations subdirectory. For example, to reproduce the first target image shown in the TOC (“target gradient from high right to low left”), go to “Interesting Patterns/Configurations/0.24124535555222237”, and then run BETSE using the top-level .yaml file in that folder (gradientVmem_Betse_config.yaml). Similarly, if the readers want to watch the whole cell tissue activity during the simulated timeline go to “Interesting Patterns/Movies/0.24124535555222237”. For selected frames from that movie the reader should go to “Interesting Patterns/Image Frames/0.24124535555222237”.
